# miR-206 Enforces a Slow Muscle Phenotype

**DOI:** 10.1101/756981

**Authors:** Kristen K. Bjorkman, Martin G. Guess, Brooke C. Harrison, Michael M. Polmear, Angela K. Peter, Leslie A. Leinwand

## Abstract

Striated muscle is a highly specialized collection of tissues with contractile properties varying according to functional needs. Although muscle fiber types are established postnatally, lifelong plasticity facilitates stimulus-dependent adaptation. Functional adaptation requires molecular adaptation, partially provided by miRNA-mediated post-transcriptional regulation. miR-206 is a muscle-specific miRNA enriched in slow muscles. We investigated whether miR-206 drives the slow muscle phenotype or is merely an outcome. We found that miR-206 expression increases in both physiologic (including female sex and endurance exercise) and pathologic conditions that promote a slow phenotype. Consistent with that observation, the slow soleus muscle of male miR-206 knockout mice displays a faster phenotype than wild-type mice. Moreover, their left ventricles have a faster myosin profile accompanied by male-specific dilation and systolic dysfunction. Thus, miR-206 appears necessary to enforce a slow skeletal and cardiac muscle phenotype and to play a key role in muscle sexual dimorphisms.

## Introduction

Skeletal muscle is a highly organized contractile tissue that composes 40% of human body mass (Janssen et al., 2000). Muscle subgroups are further specialized to perform a diverse array of functions, such as chewing, focusing and moving the eye, breathing, maintaining body posture, and both burst and sustained movements. To meet these various demands, different muscle types are characterized by different metabolic and contractile machinery. Oxidative myofibers are more fatigue-resistant and have higher mitochondrial content to facilitate β-oxidation. They can be further subdivided into slow oxidative and fast oxidative myofibers. Slow oxidative fibers are also called type I fibers due to expression of type I/β-myosin heavy chain (MyHC), a member of the sarcomeric MyHC protein family which ultimately confers contractile properties to the muscle. Fast oxidative fibers express type IIa, IIx, or a mixture of those two faster MyHCs. They also are more oxidative than type I fibers, presumably to accommodate the greater ATP demands of types IIa and IIx MyHCs compared to type I (Chemello et al., 2019). Fast glycolytic myofibers fatigue more easily and rely on glycolysis for energy production. In smaller mammals, they express the fastest MyHC, type IIb or a IIx/IIb mixture, to promote higher contraction speed while larger mammals do not express the type IIb myosin protein.

Although fiber types are established shortly after birth (Gokhin et al., 2008), adult skeletal muscle retains remarkable plasticity in order to adapt to changing demands. In addition to tuning performance, muscle is a metabolically demanding tissue that is critical to maintaining body-wide homeostasis due to the balance of energy substrate utilization. Physiological stimuli such as short sprint versus endurance exercise training shifts muscles to a more glycolytic or more oxidative phenotype, respectively (Allen et al., 2001; Andersen and Henriksson, 1977; Ross and Leveritt, 2001). Muscle unloading due to a variety of stimuli typically results in a more glycolytic profile (Caiozzo et al., 1998; Harrison et al., 2003; McCarthy et al., 1997). Certain pathologic states, including muscular dystrophy, result in a shift to more oxidative/slow fibers in part because fast fibers are more susceptible to damage (Webster et al., 1988). Finally, although understudied, evidence from rodents to humans suggests that many female muscles are slower than their male counterparts (Haizlip et al., 2015; Janssen et al., 2000).

Cardiomyocytes, the contractile cells in cardiac muscle, express two MyHCs in mammals: the same β-MyHC found in Type I skeletal muscle fibers as well as the faster, cardiac-restricted α-MyHC. There is also evidence that female cardiac muscle is slower than male (Patrizio et al., 2013; Trexler et al., 2017). While most mammalian species express both MyHCs in the heart, the α:β ratio varies between species but is tightly controlled within a species. Mice typically express >99% α-MyHC while humans express >90% β (Izumo et al., 1987; Krenz and Robbins, 2004; Miyata et al., 2000; Sadayappan et al., 2009). However, β-MyHC levels increase in most types of heart disease, regardless of species (Miyata et al., 2000; Nadal-Ginard and Mahdavi, 1989; Nakao et al., 1997). This is presumably a compensatory response as the slower, more energetically efficient β-MyHC may be more suited to the metabolic stress that typifies many forms of heart disease (Hoyer et al., 2007). However, the accompanying reduction in contractility ultimately renders this a maladaptive long-term response.

Tuning the gene expression profiles in these fiber types occurs through a coordinated transcriptional and post-transcriptional network. miRNA-mediated post-transcriptional regulation is a key event in muscle development: skeletal muscle-specific knockout of Dicer, which prevents all miRNA maturation in this tissue, is embryonic lethal (O’Rourke et al., 2007). Several miRNAs have been specifically shown to modulate muscle fiber type. For example, double knockout of the two miR-133a genes doubles the percentage of type I fibers at the expense of type II fibers in the soleus, suggesting it enforces a fast program (Liu et al., 2011). In addition, miR-27a promotes a fast oxidative (type IIa) phenotype while miR-499 and miR-208b promote the type I slow phenotype (Chemello et al., 2019; van Rooij et al., 2009). Along with the miR-208/499 and miR-133 family, the miR-1/206 family is also predominantly expressed in muscle (Ma et al., 2015; McCarthy, 2008; Mitchelson, 2015). In the basal state, miR-206 is most highly enriched in slow muscle fibers (Boettger et al., 2014; Williams et al., 2009). miR-206 is also up-regulated in fast muscle in several pathologic settings, including the chronic, progressive damage associated with amyotrophic lateral sclerosis (ALS) and Duchenne muscular dystrophy (DMD) as well as acute chemical injury from cardiotoxin injection (Liu et al., 2012; Williams et al., 2009; Yuasa et al., 2008). In conditions such as spaceflight and aging, which are associated with mixed fiber type muscle shifting towards a faster profile, miR-206 levels decrease (Allen et al., 2009; Hamrick et al., 2010; Kim et al., 2014). Notably, despite a general sex difference in muscle fiber types, it is not known whether biological sex influences miR-206 expression. Thus, we sought to determine in both sexes whether the miR-206 slow muscle enrichment is an outcome or a driver of the oxidative phenotype.

## Materials and Methods

### Cloning and mutagenesis

The minimal promoter (minTATA) and MyoG enhancer firefly luciferase reporter gene consructs were previously described (Cheung et al., 2007). We cloned the miR-206 enhancer (GRCm38/mm10 chr1:20,678,053-20,678,259) in the same manner as described for MyoG. We generated E-box point mutations (CANNTG ➔ CANNT<u>A</u>) with the QuikChange II site-directed mutagenesis kit (Agilent, 200523, Santa Clara, CA) per the manufacturer’s instructions. We verified all clones by Sanger sequencing. MRF expression constructs were a kind gift from Dr. Xuedong Liu (University of Colorado Boulder). Primer sequences are listed in Supplementary Table 1.

### Cell culture and transfection

We grew C2C12 myoblasts in Growth Medium (GM): high glucose DMEM (Invitrogen, 11960069, Waltham, MA) supplemented with 20% fetal bovine serum, 2 mM L-glutamine, 100 U/mL penicillin and 100 μg/mL streptomycin, and 1 mM sodium pyruvate. We differentiated them to myotubes by changing media to Differentiation Medium (DM): high glucose DMEM supplemented with 5% adult horse serum, 2 mM L-glutamine, 100 U/mL penicillin and 100 μg/mL streptomycin, and 1 mM sodium pyruvate. When differentiating, we refreshed DM every day to prevent media acidification. We grew 10T ½ cells in GM but with 10% FBS. For luciferase assays, we plated cells in triplicate in 6-well dishes at a density of 50,000 cells/well (C2C12) or 100,000 cells/well (10T ½) 24 hours before transfection. We repeated each experiment at least twice with independent thaws of cells. We transfected with *Trans*IT-LT1 (Mirus Bio, MIR 2305, Madison, WI) according to the manufacturer’s instructions. We co-transfected all firefly luciferase constructs with pRL-TK as a control. When differentiating C2C12 myotubes, we collected Day 0 timepoints 24 hours post-transfection and initiated differentiation at the same time for later timepoints. We collected 10T ½ cells 24 hours post-transfection. At harvest, we washed cells twice with phosphate-buffered saline (PBS) solution and stored them in an ultralow freezer in order to process all timepoints together at the end of the experiment.

### Luciferase assays

We used a Dual-Luciferase Reporter Assay System (Promega, E1960, Madison, WI) according to the manufacturer’s instructions with a Turner Designs TD-20/20 luminometer. Briefly, we lysed frozen cells in 1X Passive Lysis Buffer and measured reporter gene activities first in LARII reagent (firefly luciferase) and second in Stop & Glo reagent (Renilla luciferase). Read times for both were 10 seconds. We compared Renilla/firefly ratios.

### Mice

All wild-type C57Bl/6J (Jackson Laboratories, 000664, Sarcramento, CA), miR-206 KO (a kind gift from Dr. Eric Olson, University of Texas Southwestern Medical Center and described in (Williams et al., 2009)), and *mdx4cv* were between 4 and 7 months old. Aromatase knockout (ArKO) mice were 4 months old as this age precedes the age-dependent increase in testosterone observed in the ArKO females (Haines et al., 2012). Mice were housed in standard temperature and humidity-controlled environments with a 12-hour light-dark cycle and offered food and water *ad libidum*. We euthanized mice by cervical dislocation under inhaled isoflurane. We excised and flash froze in liquid nitrogen all tissues used for molecular analyses. For muscles used for sectioning and staining, we excised and embedded in OCT compound and then froze them in melting isopentane chilled with liquid nitrogen.

We used animals from multiple litters to achieve the pre-determined group sizes and always included littermates. We randomly allocated animals to experimental groups within a genotype. For genetically modified animals, we bred heterozygotes together to ensure littermates of both WT and genetically modified genotypes. Within a strain, we co-housed animals of the same sex to provide equivalent environments between genotypes.

### Cage Activity

To model endurance exercise, we provided a cage wheel to mice and allowed them to run voluntarily as previously described (Allen et al., 2001). To model muscle unloading, we performed hindlimb suspension as previously described (Allen et al., 2009). We suspended the tail at a 30° angle by taping it to a plastic dowel attached to a swivel apparatus. The forelimbs were still in contact with the cage bottom. We attached the suspension apparatus to a guide wire so mice were able to access all areas of the cage. We euthanized mice and harvested tissues 3, 7, or 14 days after initiating exercise or hindlimb suspension. We compared samples from exercised or hindlimb suspension mice to age-matched normal cage activity controls.

### Isoproterenol treatment

We chronically administered isoproterenol in 1 μM ascorbic acid diluted in sterile saline to C57Bl/6J wild-type male mice via subcutaneous implantation of a miniosmotic pump (model 2001; Alzet, Cupertino, CA). The pumps delivered isoproterenol at a dose of 30 or 45 mg/kg body weight per day. Vehicle control pumps contained 1 μM ascorbic acid diluted in sterile saline. We euthanized mice as described above seven days after pump implantation.

### BaCl_2_ injury

We injured the TA or gastrocnemius with barium chloride as previously described (Guess et al., 2013). Briefly, we anesthetized wild-type C57Bl/6J male mice with 1-4% inhaled isoflurane and shaved the right hind limb. We cleaned the shaved area with 70% ethanol and used a 27-gauge insulin syringe to inject the muscle with 50 μL of 1.2% BaCl_2_ dissolved in normal saline. We monitored mice for adverse effects while they recovered on a 37C heat block.

### Echo

We performed M-mode transthoracic echocardiography as previously described (Haines et al., 2012). Briefly, we mildly sedated each animal immediately before imaging with 2% inhaled isoflurane. We used a rodent-specific echodardiography machine (Visualsonics, Toronto, Canada) and a 15 MHz linear array probe to image at the midpapillary muscle level. We analyzed images with Echo-Pack software (GE Vingmed, Horten, Norway). Heart rate was the same in all groups of 206KO and WT mice (Figure 7—figure supplement 3). Each measurement per animal represents the average of three consecutive cardiac cycles.

### Hydrodynamic limb vein injection and *in vivo* imaging

We performed hydrodynamic limb vein (HLV) injections and subsequent *in vivo* reporter gene activity imaging as previously described in (Guess et al., 2013). Briefly, we co-injected the miR-206 enhancer-driven luciferase plasmid or the basal promoter-only (minTATA) control plasmid with the normalization construct p-mKate, which expresses a far red fluorescent protein from a constitutively active promoter.

### Human biopsies

Human DMD (kindly provided by Dr. Eric Hoffman of George Washington University) and healthy control vastus lateralis (VL) biopsy samples were previously described (Guess et al., 2015).

### Sectioning and immunostaining

We cut 10 μm sections with a cryostat chilled to −20C from the mid-belly of the soleus and immunostained with standard techniques. For fiber typing, primary antibodies against different MyHC isoforms were all from Developmental Studies Hybridoma Bank, Iowa City, IA (MyHC IIa (SC-71), MyHC IIx (6H1), and β-MyHC (BA-D5)). We used isotype-specific secondary antibodies (FITC-goat α-mouse IgM (Jackson 115-095-075), AlexaFluor568-goat α-mouse IgG1 (Invitrogen A-21124), and AlexaFluor647-goat α-mouse IgG2b (Jackson 15-605-207)). For cross-sectional area calculations, we delineated individual myofibers with α-laminin staining (Sigma, L-9393, St. Louis, MO) and FITC-donkey α-rabbit (Jackson 711-095-152) as secondary. We stained DNA with DAPI. We collected all images on a Nikon TE2000 inverted fluorescent microscope with a 20X objective connected to a Nikon DS-QiMc-U3 camera controlled through the NIS-Elements AR software version 4.00.03. We kept exposure times for a given channel constant for all slides. We processed raw images with the Image5D plugin in ImageJ using the same settings for a given channel across all images.

### Fiber typing

In ImageJ, we manually counted fiber types across entire soleus sections based on MyHC immunostaining. This analysis was double-blinded: before analysis, we provided images with non-descriptive, numbered labels to an investigator not involved in the study who then relabeled them and returned them for analysis. After scoring fiber types for each section, we asked for the labeling key in order to reveal genotypes.

### CSA measurements

We manually measured myofiber cross-sectional area across entire laminin-stained soleus sections using ImageJ. This analysis was also double-blinded as described in the Fiber Typing section.

### RNA isolation and qPCR analysis

We isolated total RNA with TRI Reagent (MRC, TR 118, Cincinnati, OH) according to the manufacturer’s protocol. We homogenized frozen tissue samples directly into ice cold TRI reagent using a dispersion tool (IKA, T10 Basic S1, Wilmington, NC). For pri-miR-206 analysis, we DNase treated the RNA (TURBO DNA-free kit; Thermo Fisher, AM1907, Waltham, MA) as the qPCR primers could not be intron-spanning. We performed all qPCR on an ABI 7500 Fast Real-Time PCR System. We assessed miRNA expression by TaqMan-based qPCR (Thermo Fisher, Part Number 4427975). We reverse transcribed miRNAs with a TaqMan MicroRNA Reverse Transcription Kit (Thermo Fisher, 4366596) and performed qPCR with TaqMan Universal PCR Master Mix, No AmpErase UNG (Thermo Fisher, 4324018), all according to the manufacturer’s instructions. We used the ΔΔC_T_ method to analyze relative expression. We normalized all mouse miRNAs to sno202 expression levels and human miR-206 to U6 RNA. We measured mRNA expression with SYBR Green-based qPCR. All primer sequences are listed in Supp. Table 1. We reverse transcribed total RNA with random hexamer primers using the Superscript II Reverse Transcriptase kit (Thermo Fisher, 18064-022). We performed qPCR with SYBR Green PCR Master Mix (Thermo Fisher, 4312704) according to the manufacturer’s instructions. We measured relative expression with the Pfaffl standard curve method. In regenerating TA (Fig. 2D) and isoproterenol-treated heart samples (Fig. 6A, Figure 6—figure supplement 4), we normalized RNA levels to 18S rRNA. In 206KO and WT soleus samples (Fig. 5), we normalized mRNA levels to Acta1. In the 206KO and WT heart samples (Fig. 7D and Figure 7—figure supplements 6 and 7), we normalized mRNA levels to Gapdh. Per animal, each data point is the average of technical duplicates.

### Graphing and statistical analysis

We graphed and analyzed all data for statistical significance with GraphPad Prism. We analyzed simple two-group comparisons with a Student’s t-test. We analyzed multi-group comparisons with ANOVA followed by a Bonferroni post-test. We chose a p-value of 0.05 as the significance threshold. Other statistical cutoffs are as noted in the figure legends. Data are presented as means with error bars representing standard error of the mean. The Ns for each experimental group are noted in the figure legends. To determine sample size for animal experiments, we performed a power analysis based on our preliminary data using G*Power. We assessed the presence of outliers by determining interquartile ranges and excluded data points more than 1.5X below the first or above the third quartile.

## Results

### miR-206 is associated with a slow-twitch muscle phenotype but is higher in fast female muscles

We measured miR-206 levels in three mouse lower hindlimb muscles with different fiber type compositions. In both males and females, we found a stepwise increase in miR-206 expression from the fastest muscle (tibialis anterior (TA)) to a mixed fiber type muscle (gastrocnemius and plantaris (GP)) to the slowest muscle (soleus (SOL)). Fig 1A reveals approximately 30-fold enrichment in soleus compared to TA. Interestingly, we measured significantly higher miR-206 levels in female TA compared to male (Fig. 1A, right panel), which supports our previous observation that the female TA contains a greater fraction of slow fibers than the male TA (Haizlip et al., 2015). There was no sex difference in the expression of the other myomiRs (miR-1, miR-133b, or miR-133a) in the TA (Figure 1— figure supplement 1). Since miR-206 levels can differentially respond to liganding of different estrogen receptors in breast cancer cells (Adams et al., 2007), we assessed a potential hormonal basis for the sex difference in the TA by examining miR-206 levels in female aromatase knockout (ArKO) mice, which do not produce any estrogen. In ArKO TA, we observed a significant 1.3-fold reduction in miR-206 levels (Fig. 1B), while miRs −1, −133b, and −133a were unaffected (Figure 1—figure supplement 2), suggesting a miR-206-specific phenomenon. There was no change in the ArKO GP and a downward trend of similar magnitude in the ArKO soleus (Figure 1—figure supplement 3). As it is well known that endurance exercise induces a shift towards a slow twitch phenotype (Allen et al., 2001; Andersen and Henriksson, 1977), we examined miR-206 expression in the TAs of male WT mice running voluntarily on exercise wheels. miR-206 levels increased 5.3-fold after 3 days of exercise but were not different from sedentary levels after 7 or 14 days of running (Fig. 1C). Hindlimb suspension, a model of muscle disuse that can induce a fast-twitch shift, did not result in changes in miR-206 expression (Fig. 1C).

**Figure 1.**
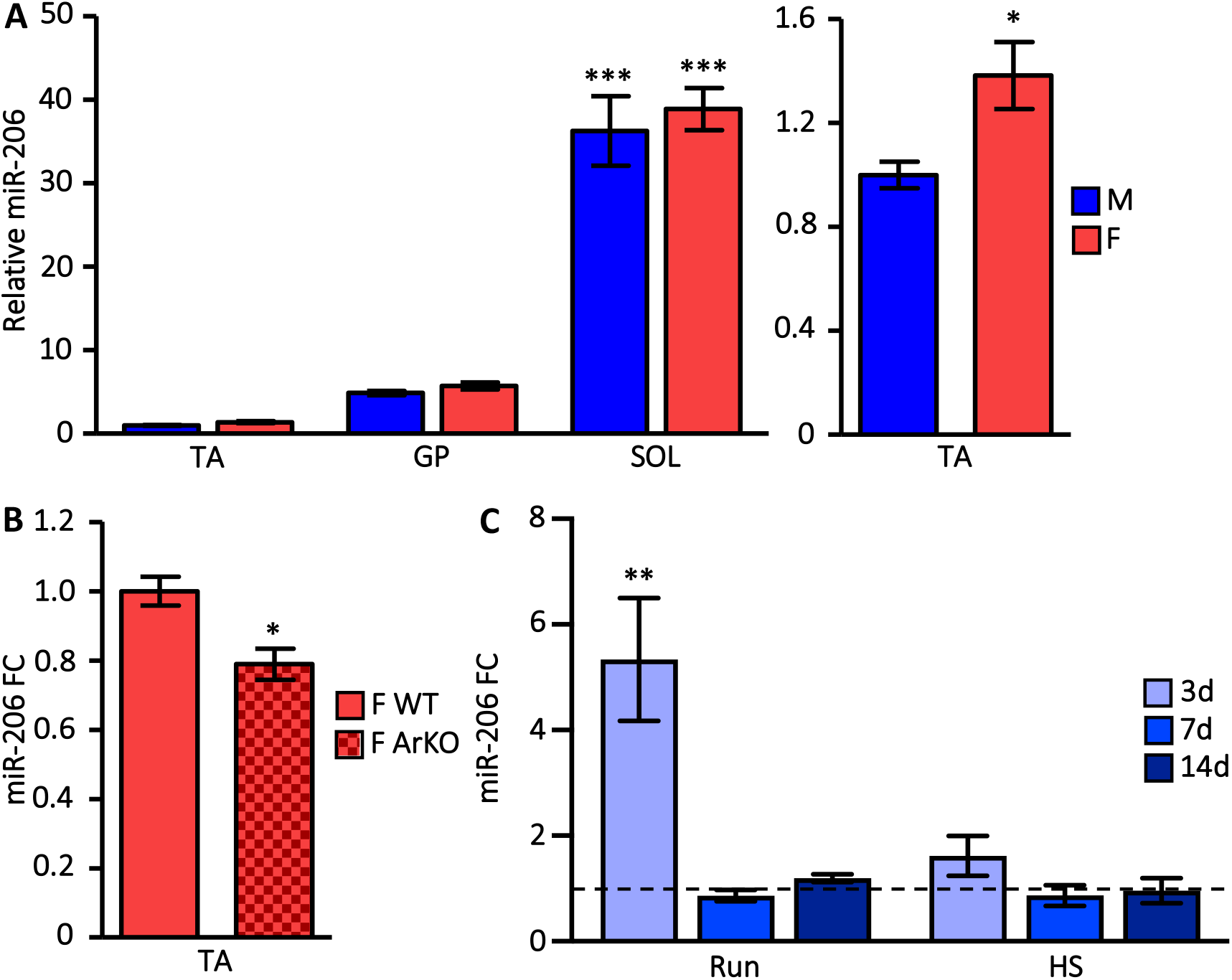
miR-206 expression is highest in slow twitch muscle and is up-regulated with endurance exercise. **(A)** miR-206 levels are highest in slow muscles in both sexes (left panel) but higher in female (F) compared to male (M) TA (right panel). We assessed miR-206 expression by qPCR in tibialis anterior (TA), gastrocnemius and plantaris (GP), and soleus (SOL) from male and female mice. Expression relative to male TA is presented. Ns are: 8 M TA, 8 F TA, 9 M GP, 8 F GP, 10 M SOL, 8 F SOL; ANOVA indicated a significant effect from muscle group (p ≤ 0.001). Asterisks indicate post-test results. *** = p ≤ 0.001 SOL vs. TA or vs. GP within each sex, * = p ≤ 0.015 F vs. M TA **(B)** miR-206 expression is lower in TAs from aromatase null compared to WT female mice. We measured miR-206 levels as in (A). Fold change (FC) relative to WT F TA is presented. N = 6 in both groups. * p = 0.014 **(C)** In male mice, miR-206 expression increased by 3 days after initiating voluntary cage wheel running and then returned to baseline but it did not change with muscle unloading by hindlimb suspension (HS). We measured miR-206 as in (A). FC relative to normal cage activity controls (represented by the dashed line at y = 1) is presented. Ns are: 6 control, 5 3d run, 3 7d run, 3 14d run, 6 3d HS, 3 7d HS, 3 14d HS; ANOVA indicated a significant effect from running (p ≤ 0.01). Asterisks indicate post-test results. ** = p ≤ 0.01 3d run vs control. 3 figure supplements accompany this figure. Source data are available in the file **Figure 1 Source Data**.

We next evaluated whether miR-206 expression responds to pathologic muscle stimuli that shift typical fast muscles to a slower phenotype. One characteristic of Duchenne muscular dystrophy (DMD) is selective sparing of slow muscle and disproportionate loss of fast muscle (Webster et al., 1988). In addition, the damage-induced regeneration in the earlier phases of the disease is accompanied by re-expression of embryonic myosin heavy chain (MyHC), one of the slowest sarcomeric myosins (Ciciliot and Schiaffino, 2010; Resnicow et al., 2010). In agreement with both of these observations, in the *mdx4cv* mouse model of DMD we saw miR-206 induced in both TA and GP with no change in the soleus (Fig. 2A), confirming previous reports as well (Liu et al., 2012). We observed this induction in human DMD vastus lateralis biopsies, a normally mixed fiber type thigh muscle (Fig. 2B). Muscle regeneration in response to acute injury also includes a transient period of slow embryonic MyHC re-expression before re-establishing the normal adult muscle phenotype. Accordingly, we found a stepwise increase in miR-206 expression during a two week regeneration time course in BaCl_2_-injured TA (Fig. 2C). This increase seems likely due to transcriptional activation as we observed a robust induction of the primary miR-206 transcript that preceded peak levels of the mature miRNA and then decreased towards basal levels as regeneration proceeded (Fig. 2D). We assessed transcriptional induction directly and *in vivo* by injecting a luciferase reporter gene controlled by a 200-bp upstream miR-206 enhancer (Figure 2—figure supplement 1) into GP that we subsequently injured with BaCl_2_. During the regeneration time course we saw a peak in luciferase activity three days post-injury, consistent with the peak in endogenous pri-miR-206 levels (Fig. 2E). This behavior was dependent on the miR-206 enhancer as there was no injury response from a construct with just a basal promoter driving luciferase (Fig. 2E and Figure 2—figure supplement 2). We also saw that this enhancer region responded to endogenous myogenic cues in differentiating C2C12 mouse myoblasts and in 10T ½ mouse fibroblasts transfected with each of the four muscle regulatory factors (MyoD, myogenin, MRF4, Myf5; Figure 2—figure supplements 3 and 4). MyoD is known to bind this region (Rao et al., 2006; Sweetman et al., 2008), and there are five E-boxes with the generic consensus CANNTG (Figure 2—figure supplement 1). A more refined muscle consensus site has been proposed that has a more specific core hexamer (CA(C/G)(C/G)TG) and extended flanking sequence (Yutzey and Konieczny, 1992). We found that the 1^st^, 2^nd^, and 5^th^ E-boxes conform well to this muscle consensus but the 3^rd^ and 4^th^ do not (Figure 2—figure supplement 5). In agreement, only the muscle consensus E-boxes are conserved in the human genome and mutation of only these E-boxes abolished the reporter response to C2C12 differentiation (Figure 2—figure supplement 6). Moreover, a reporter construct with the orthologous human region is also induced with mouse C2C12 differentiation cues, buttressing the assertion that only the three conserved E-boxes are bona fide sites of transcriptional activation (data not shown). This region has also been reported to function as a transcriptional enhancer dependent on the same E-boxes in neonatal rat cardiomyocytes (Yang et al., 2015).

**Figure 2.**
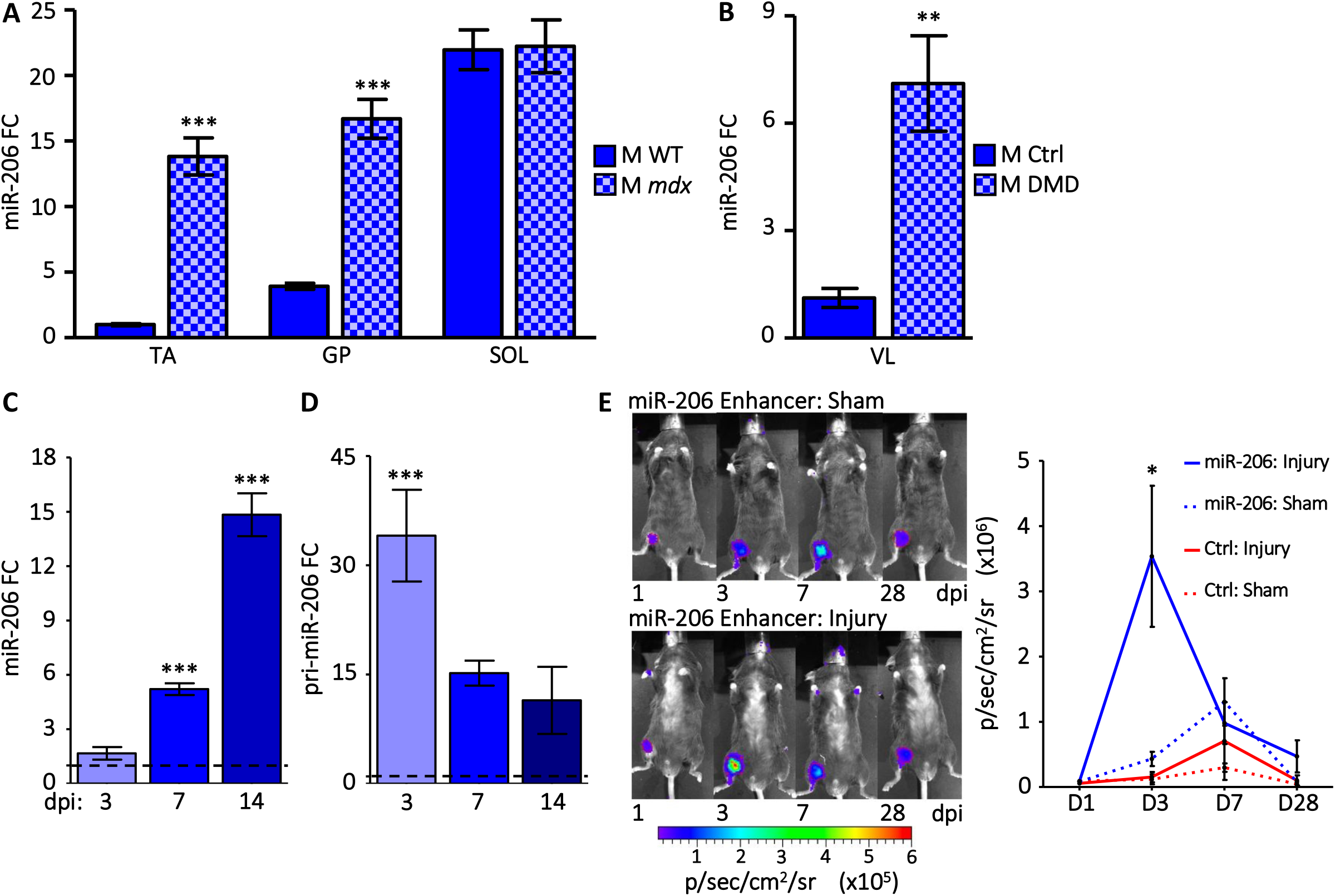
miR-206 is transcriptionally up-regulated in pathologic conditions that shift fast muscles to a slower phenotype. **(A)** miR-206 levels are higher in *mdx4cv* TA and GP compared to WT but not different in slow soleus (SOL). We measured miR-206 levels as in Figure 1. Ns are: 5 WT, 4 *mdx4cv*. ANOVA indicated a significant effect from genotype and muscle group (p ≤ 0.001). Asterisks indicate post-test results within muscle groups. *** = p ≤ 0.001 *mdx4cv* vs WT **(B)** miR-206 levels are higher in vastus lateralis (VL) from human DMD patients compared to healthy controls. We measured miR-206 levels as in Figure 1. N = 5 for both groups. ** p = 0.0090 **(C and D)** Mature miR-206 (C) and pri-miR-206 (D) levels increase in regenerating TA after BaCl_2_ injury at the indicated days post-injury (dpi). We measured mature miR-206 levels as in Figure 1. We measured pri-miR-206 by qPCR; expression levels relative to the uninjured (PBS-injected) contralateral control TA are presented. Control levels are represented as a dashed line at y = 1. Ns are: 5 3dpi, 3 7dpi, and 3 14dpi. ANOVA indicated a significant effect from injury (p ≤ 0.001). Asterisks indicate post-test results. *** = p ≤ 0.001 vs. control. **(E)** A luciferase reporter gene driven by the miR-206 enhancer is up-regulated *in vivo* during muscle regeneration. We injected the miR-206 enhancer reporter construct or the minTATA negative control reporter intravenously into the mouse hindlimb. We subsequently injured the gastrocnemius with BaCl_2_ (“injury”) or injected it with PBS as a control (“sham”). We measured reporter gene activity at days 1, 3, 7, and 28 after injury. Representative images of miR-206 reporter mice are shown. Bioluminescence signal is expressed as photons/second/cm2/steradian (p/sec/cm^2^/sr). Signal intensity was false colored according to the color bar below. Average signal intensities are plotted to the right. Ns are 3-4. * = p ≤ 0.05 vs. control group injected with the same plasmid. 6 figure supplements accompany this figure. Source data are available in the file **Figure 2 Source Data**.

### miR-206 deletion results in a slow-to-fast muscle switch

We next investigated whether miR-206 expression is an outcome or a driver of the slow muscle phenotype by examining the miR-206 knockout mouse (206KO) (Williams et al., 2009). We found a male-specific increase in soleus weight while the faster TA and GP muscles were unaffected (Fig. 3A and Figure 3—figure supplements 1 and 2). Histological examination of 206KO soleus cross-sections suggested there were larger fibers than in wild-type counterparts (representative images Fig. 3B). Indeed, we observed an enrichment for larger fibers and a depletion of small fibers in 206KO soleus when we measured cross-sectional area (Fig. 3C). In fact, only 206KO mice had any myofibers larger than 1300 μm^2^. This effect appears specific to lack of miR-206 as we did not observe any compensatory change in the closely related miR-1 nor any perturbation in the genetically linked miR-133b (Figure 3—figure supplement 3) or in linc-MD1, the non-coding RNA also generated from this locus (Figure 3—figure supplement 4). We further characterized the phenotype of the 206KO soleus through MyHC fiber typing. We stained cross-sections with antibodies specific to slow type I (β-MyHC), intermediate type IIa, and fast type IIx (representative images Fig. 4A). Relative to wild-type, we calculated a 25% decrease in the proportion of type I fibers and a 14% increase in the proportion of type IIa fibers in 206KO mice. Type IIx fibers are rare in the predominantly slow soleus, but we observed a striking increase in the fraction of these fast fibers in 206KO mice. While this was not statistically significant due to inter-animal variability, it is noteworthy (Fig. 4B). Corroborating these fiber typing data, we saw a modest change in β-MyHC mRNA but a significant 2.5-fold increase in MyHC IIa. Again, we measured the largest increase (3.4-fold) in MyHC IIx mRNA (Fig. 4C). MyHC IIb was essentially undetectable in the soleus of either genotype (data not shown).

**Figure 3.**
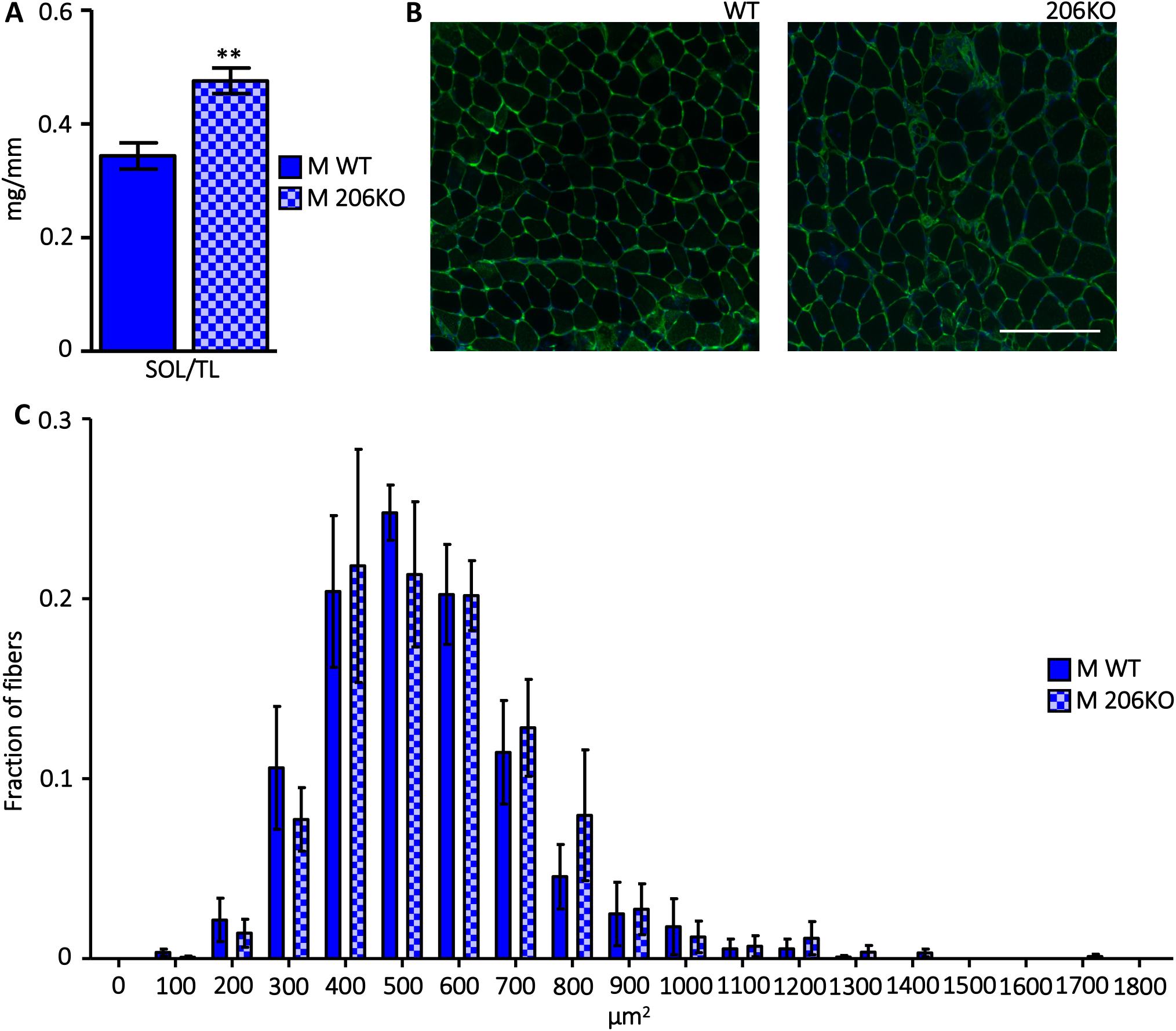
miR-206 knockout increases male soleus mass and myofiber cross-sectional area. **(A)** Soleus (SOL) mass is higher in 206KO male mice. Muscle mass normalized to tibia length (TL) is presented. Ns are 7 WT, 6 206KO; ** p = 0.0019 **(B)** Representative laminin-stained soleus cross-sections from WT (left) and 206KO (right) male mice. Scale bar denotes 100 μm. **(C)** There is a shift towards larger cross-sectional area (CSA) in myofibers from 206KO compared to WT male mice. We measured CSA across entire soleus cross-sections from 3 mice of each genotype and calculated the proportion of myofibers in 100 μm^2^ bins. 4 figure supplements accompany this figure. Source data for (A) and (C) are available in the file **Figure 3 Source Data**.

**Figure 4.**
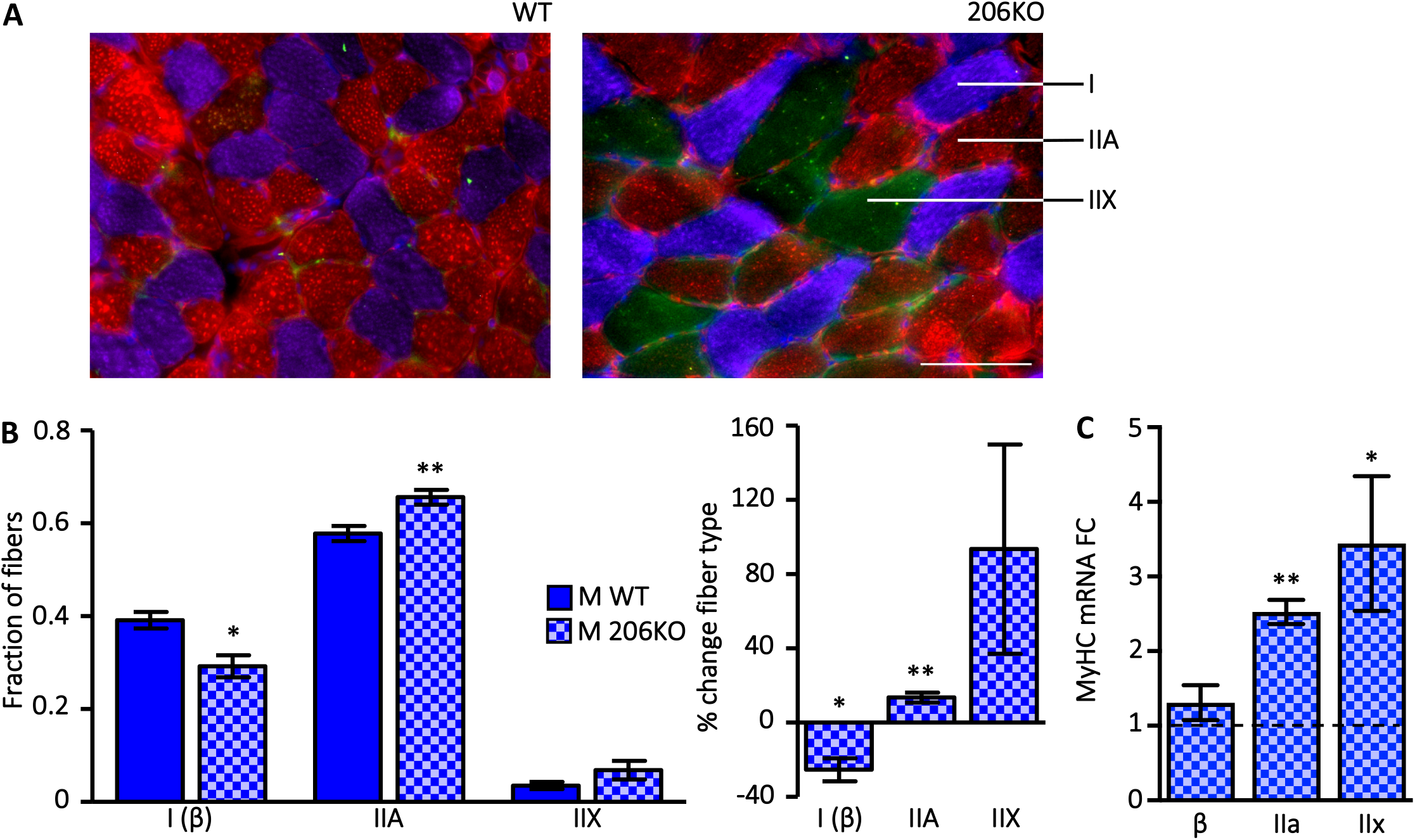
The proportion of slow fibers decreases while fast fibers increase in miR-206 KO soleus. **(A)** Representative MyHC-stained soleus cross-sections from WT (left) and 206KO (right) male mice. Type I (β-MyHC) fibers are purple, Type IIA are red, and Type IIX are green. Nuclei are revealed by DAPI staining (blue). Scale bar denotes 100 μm. **(B)** Quantification of proportions of Type I, IIA, and IIX fibers across entire soleus cross-sections from 6 WT and 4 206KO male mice (left panel) and the percent change in each fiber type (right panel; 206KO vs WT). * = p ≤ 0.05, ** = p ≤ 0.01 206KO vs WT **(C)** Fast MyHC mRNA levels increase in male 206KO soleus. We measured mRNA levels of β-MyHC, MyHC IIa, and MyHC IIx as in Fig. 2D. WT levels are represented as a dashed line at y = 1. N was 4-6 for each group. * = p ≤ 0.05, ** = p ≤ 0.01 206KO vs WT. Source data for (B) and (C) are available in the file **Figure 4 Source Data**.

To extend our molecular profiling of 206KO soleus, we measured expression levels of RNAs typically enriched in either slow or fast skeletal muscle (Fig. 5). We considered transcription factors, non-coding RNAs, troponin isoforms, and metabolic factors in addition to the MyHC signature described above. Amongst transcription factors, we found a two-fold up-regulation of the slow-associated Mef2c as well as the fast-associated Six1 and Eya1. While at first this may seem contradictory, we address this more fully in the Discussion below. We found no change in the slow-associated miRNAs miR-208b and miR-499 but a two-fold up-regulation of the fast-associated long non-coding RNA linc-MYH. Beyond the MyHC isoform content, contractile properties are also tuned by functionally distinct isoforms of other sarcomere components, including troponins. In skeletal muscle, the Tnni2 isoform of troponin I is fast-associated while Tnni1 is slow-associated. However, we did not see changes in the expression of either isoform. Expression of the slow-associated calcium-handling factor Serca2a did not change and there was a modest but significant increase in the oxygen-binding protein myoglobin (Mb). When we consider these data in conjunction with the MyHC analysis, we see an overall shift towards inducing a fast oxidative muscle phenotype in the absence of miR-206, indicating it is likely to be an enforcer of the slow program.

**Figure 5.**
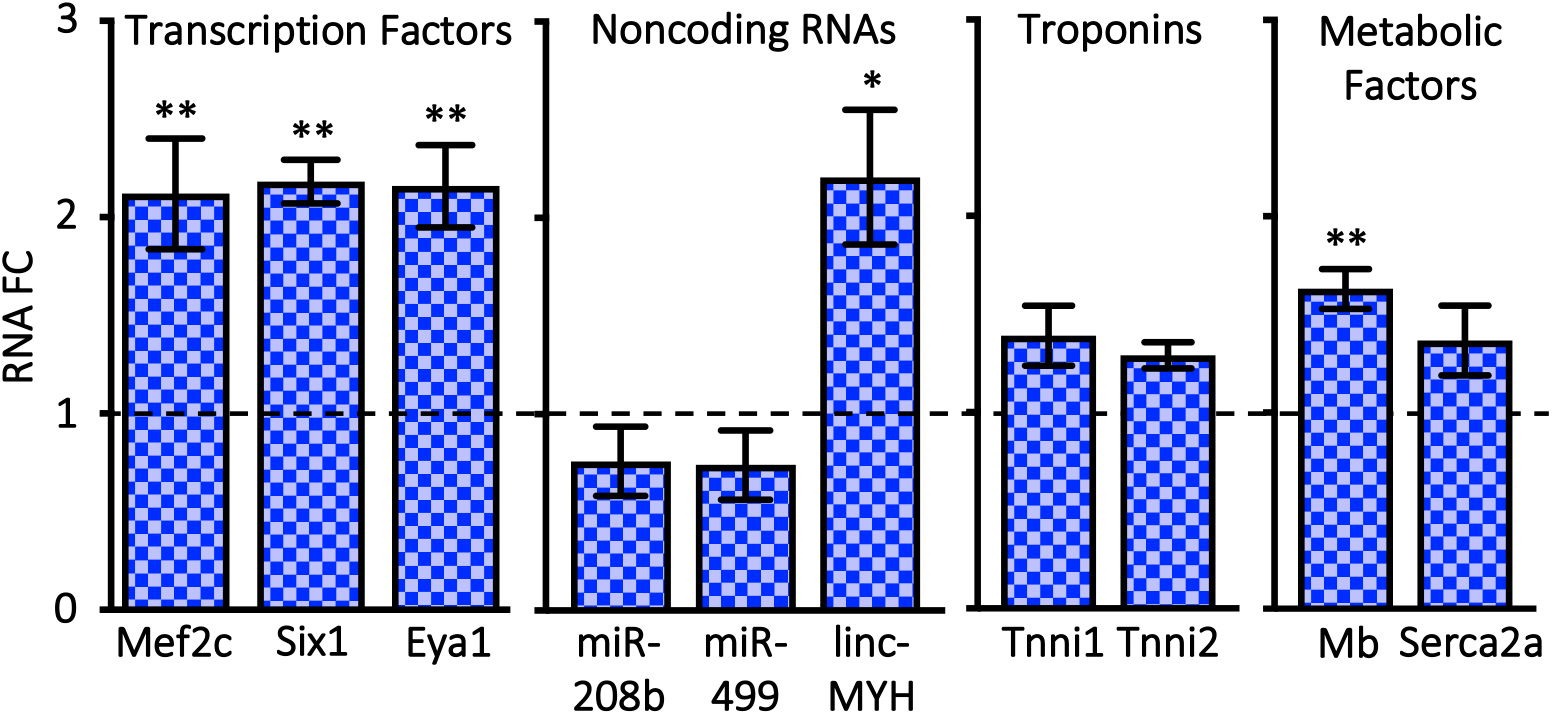
The gene expression signature of the miR-206 KO soleus shifts towards fast oxidative. We measured mRNA levels of genes from diverse functional categories in the soleus by qPCR. WT levels are represented as a dashed line at y = 1. N was 5-6. * = p ≤ 0.05, ** = p ≤ 0.01 206KO vs WT. Source data are available in the file **Figure 5 Source Data**.

### miR-206 deletion results in cardiac dilation

In a healthy rodent heart, miR-206 levels are typically quite low (Figure 6—figure supplement 1) (Boettger et al., 2014; Kim et al., 2006). However, under pathological conditions including myocardial infarction (MI), hyperglycemia, and genetic dilated cardiomyopathy, miR-206 is induced (Dong et al., 2009; Limana et al., 2011; Shan et al., 2010, 2009; Westendorp et al., 2012). Two MyHC isoforms are expressed in the heart: β-MyHC and the faster α-MyHC. One characteristic of multiple forms of heart disease is a shift towards a greater proportion of β-MyHC. To evaluate whether miR-206 induction extends to other pathologic conditions in the heart, we treated wild-type male mice with increasing doses of the β-adrenergic receptor agonist isoproterenol, which chemically models pressure overload. Confirming the pathologic impact of the treatment, we saw increases in heart rate, normalized left ventricular (LV) mass, and the mRNAs encoding the pathologic markers atrial and B-type natriuretic factors (ANF and BNP) (Figure 6—figure supplements 2, 3, and 4). We saw a significant dose-responsive increase in β-MyHC mRNA and in miR-206, the latter of which was induced four-fold in 45 mg/kg-treated animals compared to vehicle controls (Fig. 6B).

**Figure 6.**
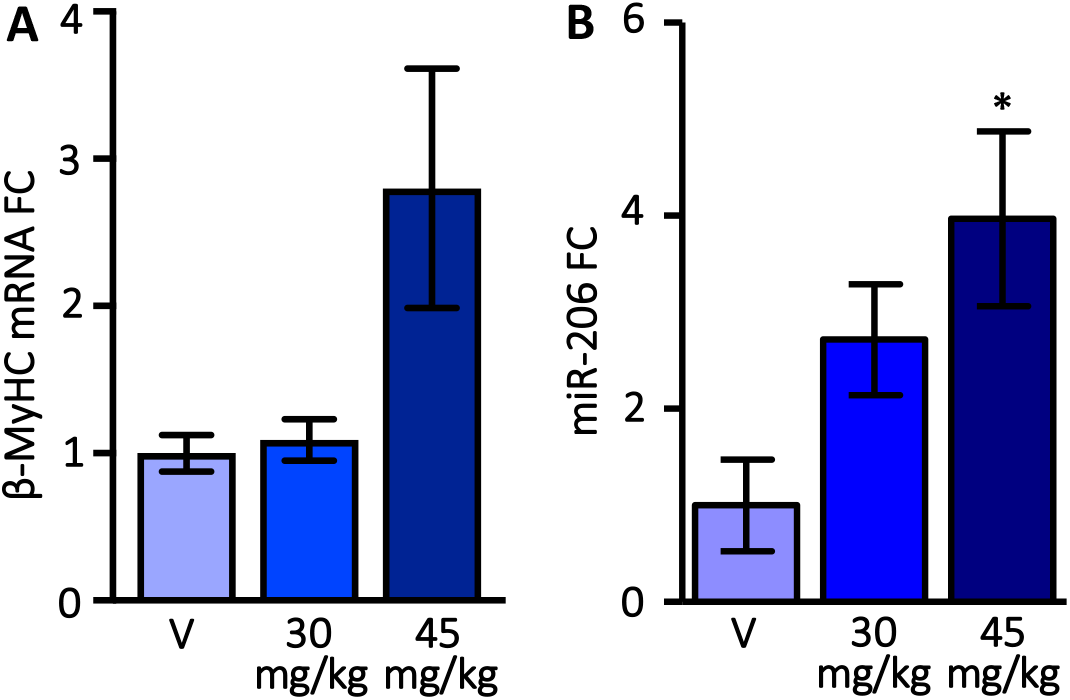
miR-206 is associated with pathologic increases in slow MyHC expression in the LV. **(A and B)** A dose-dependent increase in β-MyHC mRNA (A) and miR-206 (B) levels in the male LV with isoproterenol treatment (30 mg/kg and 45 mg/kg). We measured β-MyHC mRNA as in Fig. 2D and miR-206 as in Fig. 1. Ns are 6 for vehicle (V) and 8 for both treatments. ANOVA indicated significant treatment effects for both β-MyHC and miR-206. Asterisks indicate post-test results. * = p ≤ 0.05 vs. V. 4 figure supplements accompany this figure. Source data are available in the file **Figure 6 Source Data**.

Therefore, we proceeded to morphologically, functionally, and molecularly characterize hearts from 206KO animals. In males, we found a significant 11% increase in normalized LV mass (Fig. 7A) with no change in female LVs (Figure 7—figure supplement 1). Similar to the soleus, we saw no dysregulation of other myomiRs, including miRs-133b, −1, −208a, or −208b (Figure 7—figure supplement 2). Using M-mode echocardiography, we observed significant LV wall thinning and chamber dilation during systole in male 206KO mice compared to wild-type controls (Fig. 7B). Consistently, M-mode calculations revealed a 59% increase in male 206KO LV systolic volume and an accompanying 17% decrease in ejection fraction and 25% decrease in percent fractional shortening, which were all statistically significant (Fig. 7C). Again, we saw no change in female LV dimensions or function (Figure 7—figure supplements 4 and 5).

**Figure 7.**
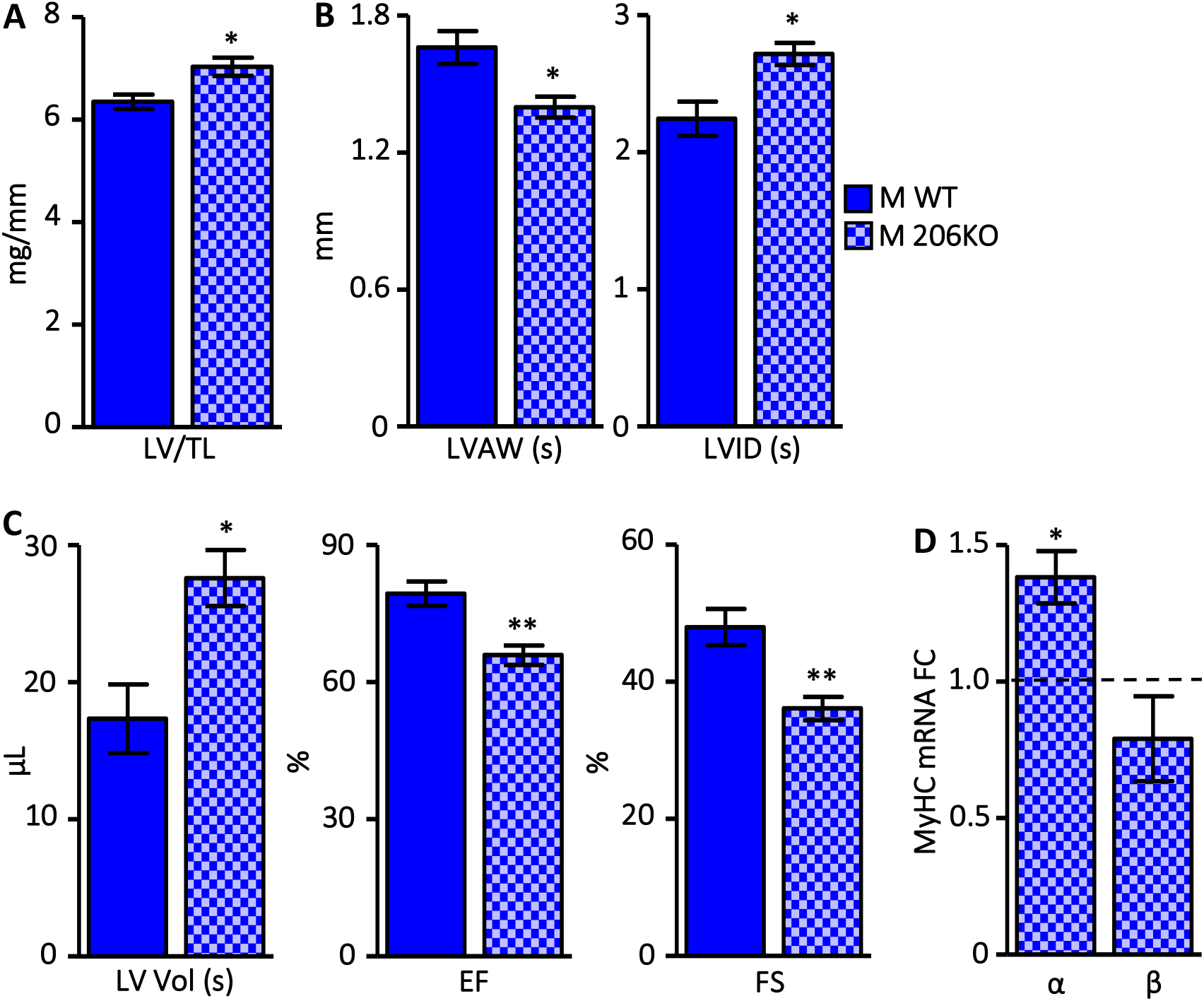
miR-206 knockout results in cardiac dilation and systolic dysfunction. **(A)** Left ventricle (LV) mass normalized to tibia length (TL) is higher in 206KO males compared to WT. Ns are 7 for WT and 6 for 206KO. * = p ≤ 0.05 vs. WT. **(B and C)** M-mode echocardiographic measurements (C) and calculations (D) indicate LV dilation and systolic dysfunction in male 206KO mice. We measured LV anterior wall (LVAW) thinning, increased LV interior diameter at systole (LVID), and increased LV volume at systole (LV Vol). Both ejection fraction (EF) and fractional shortening (FS) decreased. Ns are 4 for WT and 5 for 206KO. * = p ≤ 0.05, ** = p ≤ 0.01 206KO vs. WT. **(D)** mRNA corresponding to the faster cardiac α-MyHC increases while the slower β-MyHC decreases in male 206KO LV compared to WT. We measured MyHC mRNA levels by qPCR as in Fig. 2D. WT levels are represented as a dashed line at y = 1. Ns are 5-6. * = p ≤ 0.05 206KO vs WT. 7 figure supplements accompany this figure. Source data are available in the file **Figure 7 Source Data**.

We molecularly assessed the male 206KO LVs by qPCR-based analysis of a suite of mRNAs and miRNAs whose expression is frequently perturbed in a variety of cardiac pathologies (Taegtmeyer et al., 2010; Van Rooij et al., 2006). These include mRNAs encoding ANF and BNP, the skeletal muscle isoform of sarcomeric actin (Acta1), the calcium-handling factors Serca2a and Pln, the fetal cardiac transcription factors Gata4 and Nkx2.5, the Nfat target myocyte-enriched calcineurin-interacting protein (mCip1.4), the pro-fibrotic collagen 1 (Col1a1), and the anti-hypertrophic muscle cytokine myostatin (Mstn). Surprisingly, the only changes we observed were a 1.7-fold increase in Acta1 and a 1.6-fold decrease in Mstn, consistent with cardiac pathology and an increase in LV mass, respectively (Figure 7—figure supplement 6). We also observed no change in 10 miRNAs previously identified as part of a signature set of miRNAs whose expression levels respond to three different pathologic cardiac stimuli (Figure 7—figure supplement 7) (Van Rooij et al., 2006). However, when we measured MyHC mRNA levels, we saw a significant 1.4-fold induction of the faster cardiac isoform α-MyHC with a 1.3-fold downward trend in the slower β-MyHC (Fig. 7D). This suggests a shift towards a faster cardiac phenotype with pathologic functional consequences in the absence of miR-206 expression.

## Discussion

We show that miR-206 is essential to enforce a slow muscle program in both skeletal muscle and the heart. In healthy skeletal muscle, miR-206 expression is strongly associated with a slow oxidative phenotype and is actually significantly higher in the female compared to male TA (Fig. 1A). This aligns with our previous observations that the female TA contains a greater proportion of slow fibers and that females have a greater, but hormone-dependent, capacity for endurance exercise (Haines et al., 2012; Haizlip et al., 2015). There is likely to be a hormonal component to this expression pattern as aromatase null females express less miR-206 in the TA than wild-type females (Fig. 1B). Interestingly, estrogen can regulate miR-206 expression levels in breast cancer cells, further reinforcing this hormonal link (Adams et al., 2007). We did not see the same sex bias or response to aromatase deletion in expression of the miR-206 family member miR-1 nor in the genetically linked miR-133b, suggesting a specialized role for miR-206 in promoting the slower female phenotype.

The same miR-206 KO mouse studied here was shown to have delayed muscle regeneration after cardiotoxin injury, impaired satellite cell differentiation, and delayed neuromuscular reinnervation (Liu et al., 2012; Williams et al., 2009). Curiously, a separate knockout mouse that is lacking not only miR-206 but also miR-133b and a portion of the muscle regulatory long noncoding RNA linc-MD1 (including the sequence encoding a miR-133 sponge) displayed no overt phenotype at baseline, after acute muscle injury, or in the context of muscular dystrophy (Boettger et al., 2014). However, miR-206 and miR-133 family members have been shown to play opposing roles, where the former is pro-differentiation and anti-proliferative while the latter is pro-proliferative (Chen et al., 2006; Kim et al., 2006). Moreover, double knockout of miR-133 family members miR-133a-1 and miR-133a-2 (which, collectively, differ only at the terminal 3’ position and so are presumed to target overlapping sets of transcripts) results in an increase in oxidative myofibers (Liu et al., 2011). This also supports opposing roles for miR-206 and the miR-133 family when considered with the increased proportion of fast fiber types in male 206KO soleus that we observed (Fig. 4), potentially complicating interpretation of the miR-206/miR-133b double knockout mouse. Supporting these observations, in Fig. 5 we show higher levels of the key fast muscle transcription factors Six1 and Eya1 (Grifone et al., 2004) as well as the fast-associated lncRNA linc-MYH, which is encoded in the fast MyHC genomic locus, shares a Six1/Eya1-dependent enhancer with these MyHCs, and promotes fast-type gene expression (Sakakibara et al., 2014). The Six1 3’ UTR harbors a predicted target site for another slow muscle-specific miRNA family, miR-208/499 (TargetScan.org). Transgenic miR-499 overexpression has been shown to convert all fast fibers in the mouse soleus to slow fibers (van Rooij et al., 2009). We did not observe differential expression of either miRNA in the miR-206 KO soleus, suggesting that miR-206 could be acting downstream of miR-499 and miR-208b in promoting the slow fiber program. We also saw increased Mef2c and myoglobin mRNA levels in the miR-206 KO soleus (Fig. 5), which may initially seem to contradict a fast-type phenotypic switch as expression of these genes is associated with an oxidative profile. However, a recent gene expression analysis of single soleus myofibers revealed that fast oxidative fibers (Types IIa and IIx) are characterized by a stronger mitochondrial and electron transport chain signature than Type I, which is consistent with our overall change in gene expression and fiber type profiles (Chemello et al., 2019).

miR-206 is well-studied in the context of the early stages of myogenesis, particularly in immortalized and primary myoblast cell culture models where it inhibits proliferation and promotes the transition to differentiation by targeting transcripts such as *Pola1*, *Pax7*, and *Pax3* (reviewed in (McCarthy, 2008; Mitchelson, 2015)). Although the targeting network through which it enforces a slow muscle phenotype is not yet clear, it notably targets the class II histone deacetylase (HDAC) *Hdac4* through translational inhibition (Williams et al., 2009; Winbanks et al., 2011). Hdac4 inhibits slow muscle gene expression and, when knocked out in mice along with other class II HDACs, results in an increased proportion of Type I myofibers (Potthoff et al., 2007). This targeting relationship was established in the same miR-206 KO mice studied here when they were examined in the context of ALS (Williams et al., 2009).

In the heart, we also found that miR-206 promotes a slow phenotype. Although rodents express very little β-MyHC in the ventricles, these levels are not inconsequential and there are distinct regions where β-MyHC predominates (Krenz et al., 2007). In addition, the α-MyHC:β-MyHC ratio is strictly regulated. Interestingly, β-MyHC expression is reportedly four-fold higher in the left ventricles of sexually mature female mice compared to males, and this difference is hormone-dependent (Patrizio et al., 2013). We have also previously reported that isolated female rat cardiomyocytes are slower to reach both peak shortening and relaxation (Trexler et al., 2017). These observations are consistent with our comparison of female and male TA muscle. As miR-206 levels are very low in the healthy ventricle, we could not reliably compare male and female expression levels. However, if miR-206 is also an enforcer of slow gene expression in the heart, we predicted that: 1) in the absence of miR-206, the heart would shift towards a faster phenotype and 2) miR-206 expression would increase in pathologic conditions when the heart shifts towards a slower phenotype. Accordingly, we observed a significant increase in α-MyHC expression and a downward trend in β-MyHC expression in miR-206 KO LVs (Fig. 7D). Supporting the assertion that the small amount of β-MyHC that is expressed in the mouse heart is still critical for normal function, we observed ventricular dilation and systolic dysfunction in miR-206 KO mice (Figs. 7B, C). As in skeletal muscle, this is a male-specific phenomenon, the mechanistic basis for which is an intriguing area for future investigation. Interestingly, transgenic mice overexpressing miR-206 experience cardiac hypertrophy which, combined with our knockout data, suggests the importance of maintaining miR-206 levels within a narrow range at baseline (Yang et al., 2015). However, miR-1 levels also increased in these transgenic miR-206 mice, and miR-1 overexpression in the heart is known to be detrimental (Zhao et al., 2005).

In support of our second prediction, we observed a dose-dependent increase in miR-206 expression in the left ventricles of mice treated with the β-adrenergic receptor agonist isoproterenol. Several reports have also noted increased miR-206 expression after MI in multiple animal models (Dong et al., 2009; Limana et al., 2011; Shan et al., 2009). Both MI and β-adrenergic stimulation are associated with an increase in β-MyHC expression at the expense of the faster α-MyHC ((Harada et al., 1999); Fig. 6A). This may be a compensatory acute stress-adaptation mechanism as β-MyHC is a more energetically efficient motor. We speculate that miR-206 KO mice would be less able to adapt to cardiac stress as they may not properly induce β-MyHC. Indeed, transgenic mice expressing a miR-206 sponge develop larger infarcts after ischemia/reperfusion injury than wild-type animals (Yang et al., 2015). As this miRNA sponge may also inhibit miR-1, which is the most abundant cardiac miRNA, it will be important to examine the response of miR-206 KO mice to cardiac injury in the future.

Collectively, our data indicate that miR-206 enforces slow muscle gene expression, which ultimately tunes both skeletal and cardiac muscle performance. Moreover, the consistent protective role of miR-206 under pathologic conditions in both tissues makes it an attractive target for RNA-based therapeutics.

## Acknowledgments

We thank Dr. Eric Olson (UT-Southwestern) for generously providing the miR-206 KO mice. We also thank Amy Perry for performing echocardiography measurements, Darrian Bugge and Dr. Alberto Rossi for technical assistance, Drs. Emily Pugach, Christa Trexler, and Massimo Buvoli for valuable conversations and insight, Dr. Sarah Lehman for critical reading and feedback, and the University of Colorado Boulder BioFrontiers Advanced Light Microscopy Core facility for support with fluorescence microscopy.

## Figure Legends

**Figure 1-figure supplement 1. Expression of miR-1, miR-133b, and miR-133a is not different between male and female mice in the TA.** We measured miRNA levels by qPCR as in Fig. 1. Ns are 7-8. Source data are available in the file **Figure 1 Source Data**.

**Figure 1-figure supplement 2. miR-206 expression is not different between WT and aromatase KO (ArKO) females in the gastrocnemius and plantaris (GP) and the soleus (SOL).** We measured miRNA levels by qPCR as in Fig. 1. Ns are 5-6. Source data are available in the file **Figure 1 Source Data.**

**Figure 1-figure supplement 3. Expression of miR-1, miR-133b, and miR-133a is not different between WT and ArKO female mice in the TA.** We measured miRNA levels by qPCR as in Fig. 1. Ns are 5-6. Source data are available in the file **Figure 1 Source Data.**

**Figure 2-figure supplement 1. A 207-bp region approximately 1 kb upstream of the miR-206 locus contains spikes in conservation corresponding to E-box consensus sites.** The line drawing below the image (image from Genome Browser (genome.ucsc.edu); blue histogram illustrates conservation across placental mammals) indicates the location of the E-boxes conforming to the consensus sequence CANNTG, with E-boxes conserved in human, mouse, and rat shown in red and those not conserved in human shown in cyan. The complete mouse sequence cloned for reporter gene analysis is also shown with E-boxes similarly color-coded.

**Figure 2-figure supplement 2. *In vivo* imaging of basal promoter-driven reporter gene expression.** Reporter gene activity imaging of minTATA negative control reporter as described and quantified in Fig. 2E. Signal intensity was false colored according to the color bar below.

**Figure 2-figure supplement 3. Activity of a firefly luciferase reporter gene with the 200-bp genomic region cloned upstream of a minimal TATA-box-containing promoter increases during differentiation of C2C12 cells.** We transfected C2C12 cells with the constructs indicated on the x-axis. We induced differentiation 24 hours post-transfection and measured reporter activity at Day 0 (proliferating myoblasts), Day 1, and Day 4 after differentiation initiation. MyoG serves as a positive control and has the myogenin enhancer driving firefly luciferase. We performed all experiments in triplicate. 1-way ANOVA indicated a significant effect from day of differentiation (p ≤ 0.01). Asterisks indicate post-test results. ** = p ≤ 0.01 vs. Day 0, double daggers = p ≤ 0.01 vs. Day 1. Source data are available in the file **Figure 2 Source Data**.

**Figure 2-figure supplement 4. The miR-206 200-bp reporter responds to expression of muscle regulatory factors (MRFs) in mouse 10T1/2 fibroblasts.** We transfected 10T1/2 cells with the reporters indicated on the x-axis (described in (Figure 2-figure supplement 2)) and either empty vector control or overexpression constructs for Flag-tagged MRFs. MyoG and miR-206 200-bp both respond to all 4 MRFs, but the strength of activation in the presence of the various MRFs differs between the reporters. We performed all experiments in triplicate. 1-way ANOVA indicated a significant effect from MRF expression (p ≤ 0.0001). Asterisks indicate post-test results vs Empty Vector. ** = p ≤ 0.01; *** = p ≤ 0.001. Source data are available in the file **Figure 2 Source Data**.

**Figure 2-figure supplement 5. The miR-206 transcriptional enhancer contains 3 muscle-specific conserved E-boxes.** We analyzed E-boxes from the mouse 200-bp region (numbered from the 5’ to the 3’ end of the cloned region as cartooned in Figure 2-figure supplement 1) and the orthologous human region for conformation to the longer muscle-specific E-box consensus sequence described by Buskin and Hauschka. The consensus is above the table with the core E-box in red text. The scoring rubric is 0 points for divergence, 0.5 points for a degenerate sequence match, and 1 point for an invariant sequence match (8 = highest possible score). Asterisks indicate divergence from the muscle consensus sequence. Ø indicates that there is no orthologous human CANNTG sequence.

**Figure 2-figure supplement 6. Only conserved E-boxes contribute to transcriptional activation from the miR-206 enhancer.** We mutated E-boxes in the WT miR-206 200-bp construct from CANNTG to CANNTA. We mutated conserved E-boxes 1, 2, and 5 individually while we mutated unconserved E-boxes 3 and 4 together in one construct. Finally, we mutated all 5 E-boxes in the construct named E-boxless. We transfected C2C12 cells with each mutant construct as well as the WT and allowed them to differentiate for one day. We normalized firefly luciferase activity to control renilla luciferase activity. 2-way ANOVA indicated significant effects from reporter and day of differentiation (p ≤ 0.0001). Asterisks indicate post-test results vs D0. *** = p ≤ 0.001. We performed all experiments in triplicate. Source data are available in the file **Figure 2 Source Data**.

**Figure 3-figure supplement 1. Gastrocnemius and plantaris (GP) and tibialis anterior (TA) muscle masses are not different between 206KO and WT male mice.** We normalized muscle masses to tibia length (TL). Ns are: 7 WT GP, 6 206KO GP, 7 WT TA, 5 206KO TA. Source data are available in the file **Figure 3 Source Data**.

**Figure 3-figure supplement 2. Soleus (SOL), gastrocnemius and plantaris (GP), and tibialis anterior (TA) muscle masses are not different between 206KO and WT female mice.** We normalized muscle masses to tibia length (TL). Ns are 5 for WT, 3 for 206KO. Source data are available in the file **Figure 3 Source Data**.

**Figure 3-figure supplement 3. We verified that miR-206 is undetectable in the 206KO male soleus and found no compensatory change in miR-1 or miR-133b expression.** We measured miRNA expression as in Fig. 1. Ns are 6 for both genotypes. *** = p ≤ 0.001 vs. WT. Source data are available in the file **Figure 3 Source Data**.

**Figure 3-figure supplement 4. linc-MD1 expression is not different between male 206KO and WT soleus.** We measured RNA expression as in Fig. 5. Ns are 7 for WT, 6 for 206KO. Source data are available in the file **Figure 3 Source Data**.

**Figure 6-figure supplement 1. miR-206 expression is low in the healthy LV.** We measured miR-206 expression in the male mouse LV and TA and found expression to be over 3 orders of magnitude higher in the TA. We measured miRNA expression as in Fig. 1. Ns are 6 for the LV and 8 for the TA. *** p = 1.11×10^-9^ vs. LV. Source data are available in the file **Figure 6 Source Data**.

**Figure 6-figure supplement 2. Heart rate significantly increased after 7 days treatment with both 30 mg/kg and 45 mg/kg isoproterenol.** We measured heart rate in all animals before (solid bars) and after (patterened bars) treatment. ** = p ≤ 0.01; *** = p ≤ 0.001 vs. pre-treatment. Source data are available in the file **Figure 6 Source Data**.

**Figure 6-figure supplement 3. Left ventricle (LV) mass is higher in both groups of isoproterenol-treated animals compared to vehicle controls.** We measured LV mass and normalized to tibia length (TL). ANOVA indicated a significant treatment effect. *** = p ≤ 0.001 vs. vehicle. Source data are available in the file **Figure 6 Source Data**.

**Figure 6-figure supplement 4. Gene expression of β-MyHC, ANF, and BNP in the LVs of isoproterenol-treated mice.** We measured mRNA levels as in Fig. 2D. ANOVA indicated a significant treatment effect. Asterisks indicate post-test results. * = p ≤ 0.05; ** = p ≤ 0.01 vs. vehicle. Ns are 6 for vehicle control and 8 for both 30 mg/kg and 45 mg/kg for all measurements. Source data are available in the file **Figure 6 Source Data**.

**Figure 7-figure supplement 1. Left ventricle (LV) mass is not different between 206KO females and WT.** We measured LV mass and normalized to tibia length (TL). N was 5 for WT and 3 for 206KO. Source data are available in the file **Figure 7 Source Data**.

**Figure 7-figure supplement 2. myomiR expression in miR-206 KO vs WT left ventricles (LVs).** There is no difference in expression of miRs −133b, −1, −208a, or −208b in the male 206KO LV compared to WT. We measured miRNA expression as in Fig. 1. Ns are 4-6. Source data are available in the file **Figure 7 Source Data**.

**Figure 7-figure supplement 3. Heart rate was the same between miR-206 KO and WT males and females.** Ns were 4 for WT male (M), 5 for 206KO M, 5 for WT female (F), and 4 for 206KO F. Source data are available in the file **Figure 7 Source Data**.

**Figure 7-figure supplement 4. M-mode echocardiographic measurements revealed no changes in LV dimensions in female 206KO mice compared to WT.** LV anterior wall at systole = LVAW (s), LV interior diameter at systole = LVID (s). Ns are 5 for WT and 4 for 206KO. Source data are available in the file **Figure 7 Source Data**.

**Figure 7-figure supplement 5. M-mode echocardiographic calculations revealed no changes in LV function in female 206KO mice compared to WT.** LV volume at systole = LV Vol (s), ejection fraction = EF, and fractional shortening = FS. Ns are 5 for WT and 4 for 206KO. Source data are available in the file **Figure 7 Source Data**.

**Figure 7-figure supplement 6. Expression of mRNA cardiac stress markers in male miR-206 KO vs WT LVs.** We measured mRNA levels corresponding to the indicated genes as in Fig. 6. Acta1 was significantly up-regulated while Mstn was significantly down-regulated. WT levels are represented as a dashed line at y = 1. Ns are 5-7. * = p ≤ 0.05 206KO vs WT. Source data are available in the file **Figure 7 Source Data**.

**Figure 7-figure supplement 7. Expression of miRNA cardiac stress markers in male miR-206 KO vs WT LVs.** We measured miRNA levels corresponding to the indicated genes as in Fig. 1. WT levels are represented as a dashed line at y = 1. Ns are 7 for WT and 6 for 206KO. Source data are available in the file **Figure 7 Source Data**.

**Supplementary Table 1.**
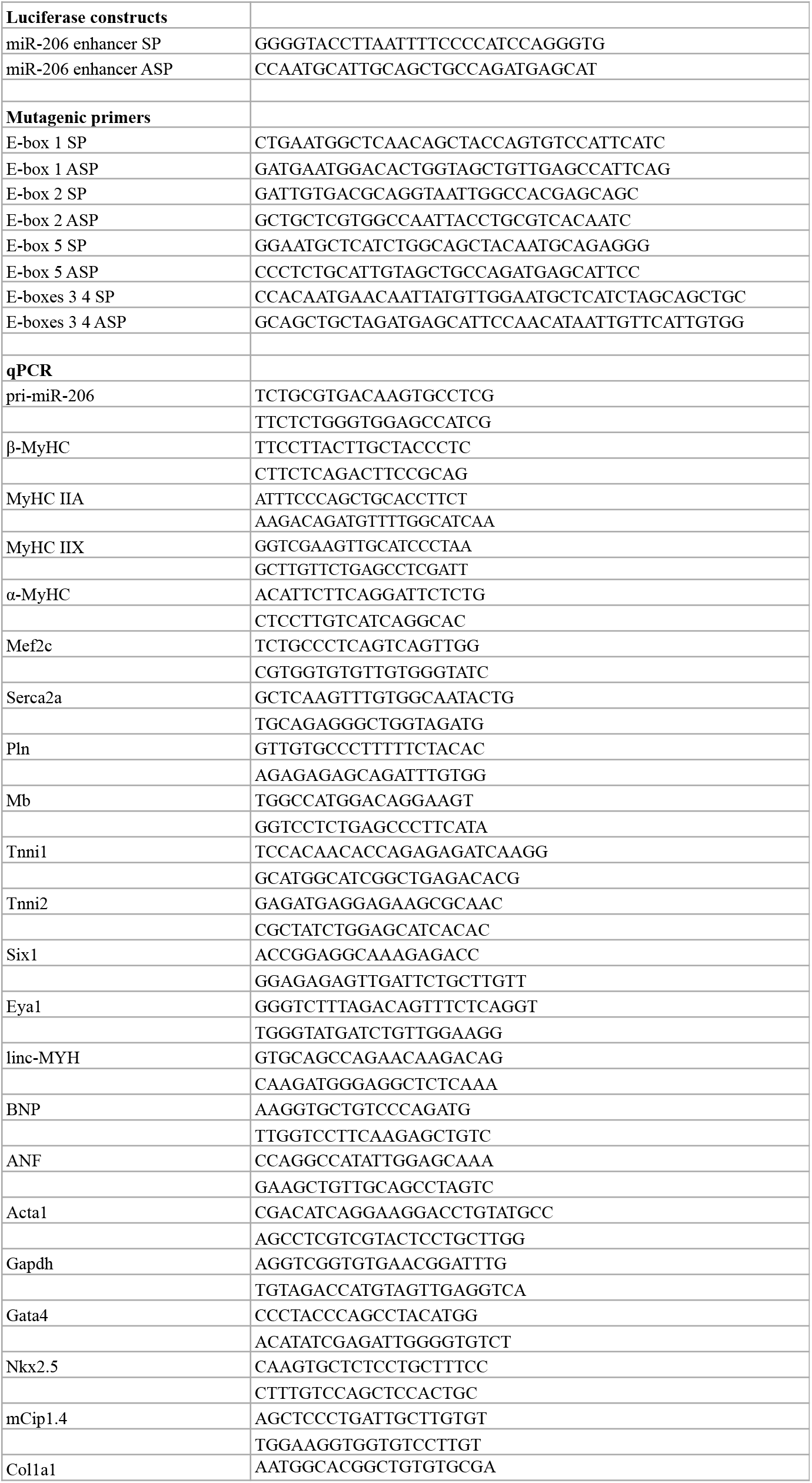
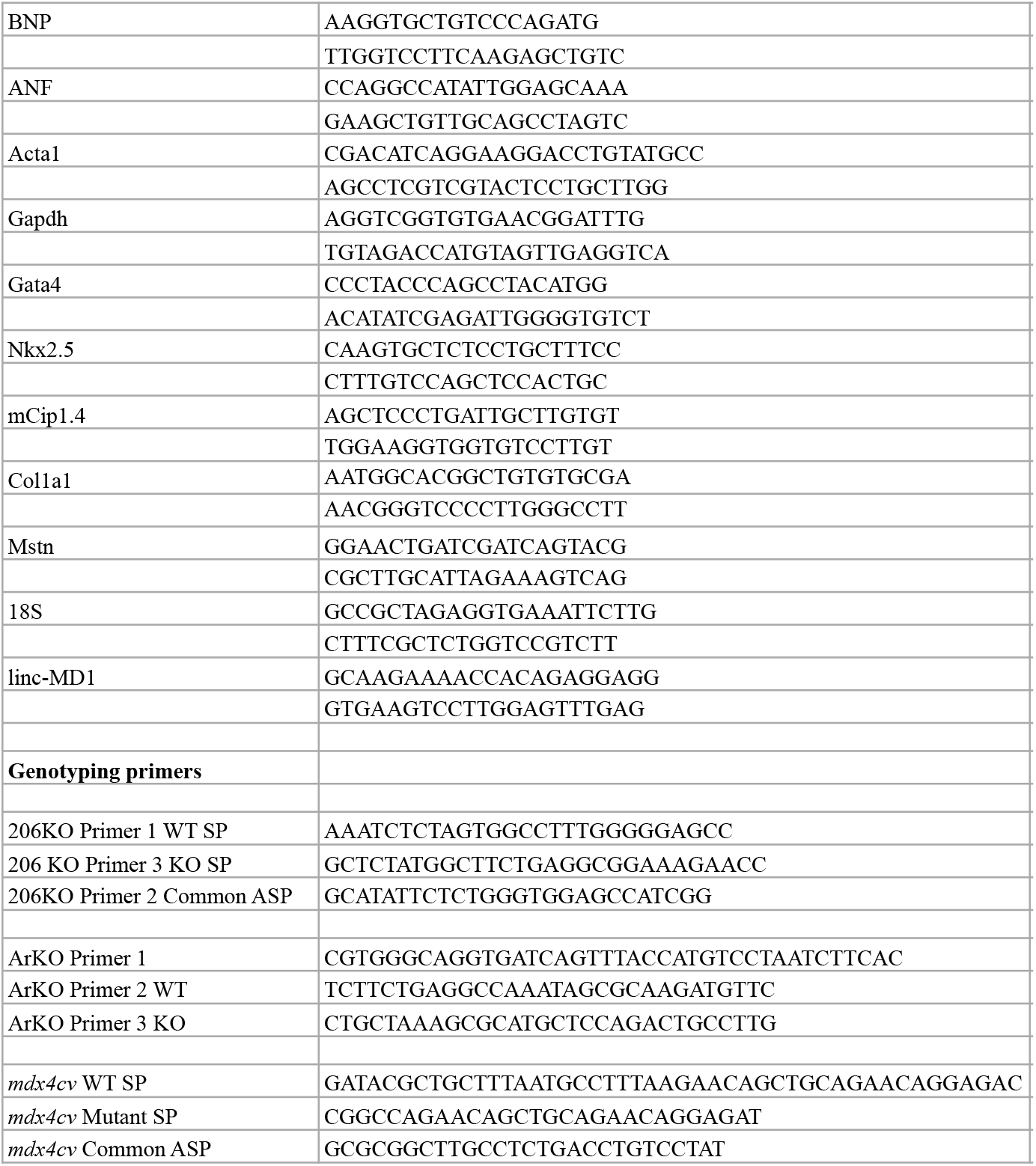
Primer sequences. Primers for cloning the miR-206 enhancer, mutagenizing the E-boxes in the miR-206 enhancer, qPCR analysis, and genotyping mice are presented 5’ ➔ 3’.

## References

Adams BD, Furneaux H, White BA. 2007. The micro-ribonucleic acid (miRNA) miR-206 targets the human estrogen receptor-α (ERα) and represses ERα messenger RNA and protein expression in breast cancer cell lines. Mol Endocrinol 21:1132–1147. doi:10.1210/me.2007-0022

Allen DL, Bandstra ER, Harrison BC, Thorng S, Stodieck LS, Kostenuik PJ, Morony S, Lacey DL, Hammond TG, Leinwand LL, Argraves WS, Bateman TA, Barth JL. 2009. Effects of spaceflight on murine skeletal muscle gene expression. J Appl Physiol 106:582–595. doi:10.1152/japplphysiol.90780.2008

Allen DL, Harrison BC, Maass A, Bell ML, Byrnes WC, Leinwand LA. 2001. Cardiac and skeletal muscle adaptations to voluntary wheel running in the mouse. J Appl Physiol 90:1900–1908.

Andersen P, Henriksson J. 1977. Training Induced Changes in the Subgroups of Human Type II Skeletal Muscle Fibres. Acta Physiol Scand 99:123–125. doi:10.1111/j.1748-1716.1977.tb10361.x

Boettger T, Wüst S, Nolte H, Braun T. 2014. The miR-206/133b cluster is dispensable for development, survival and regeneration of skeletal muscle. Skelet Muscle 4:23. doi:10.1186/s13395-014-0023-5

Caiozzo VJ, Baker MJ, Baldwin KM. 1998. Novel transitions in MHC isoforms: Separate and combined effects of thyroid hormone and mechanical unloading. J Appl Physiol 85:2237–2248. doi:10.1152/jappl.1998.85.6.2237

Chemello F, Grespi F, Zulian A, Cancellara P, Hebert-Chatelain E, Martini P, Bean C, Alessio E, Buson L, Bazzega M, Armani A, Sandri M, Ferrazza R, Laveder P, Guella G, Reggiani C, Romualdi C, Bernardi P, Scorrano L, Cagnin S, Lanfranchi G. 2019. Transcriptomic Analysis of Single Isolated Myofibers Identifies miR-27a-3p and miR-142-3p as Regulators of Metabolism in Skeletal Muscle. Cell Rep 26:3784–3797. doi:10.1016/j.celrep.2019.02.105

Chen JF, Mandel EM, Thomson JM, Wu Q, Callis TE, Hammond SM, Conlon FL, Wang DZ. 2006. The role of microRNA-1 and microRNA-133 in skeletal muscle proliferation and differentiation. Nat Genet 38:228–233. doi:10.1038/ng1725

Cheung TH, Barthel KKB, Yin LK, Liu X. 2007. Identifying pattern-defined regulatory islands in mammalian genomes. Proc Natl Acad Sci U S A 104:10116–10121. doi:10.1073/pnas.0704028104

Ciciliot S, Schiaffino S. 2010. Regeneration of Mammalian Skeletal Muscle: Basic Mechanisms and Clinical Implications. Curr Pharm Des 16:906–914. doi:10.2174/138161210790883453

Dong S, Cheng Y, Yang J, Li J, Liu X, Wang X, Wang D, Krall TJ, Delphin ES, Zhang C. 2009. MicroRNA expression signature and the role of MicroRNA-21 in the early phase of acute myocardial infarction. J Biol Chem 284:29514–29525. doi:10.1074/jbc.M109.027896

Gokhin DS, Ward SR, Bremner SN, Lieber RL. 2008. Quantitative analysis of neonatal skeletal muscle functional improvement in the mouse. J Exp Biol 211:837–843. doi:10.1242/jeb.014340

Grifone R, Laclef C, Spitz F, Lopez S, Demignon J, Guidotti J-E, Kawakami K, Xu P-X, Kelly R, Petrof BJ, Daegelen D, Concordet J-P, Maire P. 2004. Six1 and Eya1 Expression Can Reprogram Adult Muscle from the Slow-Twitch Phenotype into the Fast-Twitch Phenotype. Mol Cell Biol 24:6253–6267. doi:10.1128/mcb.24.14.6253-6267.2004

Guess MG, Barthel KKB, Harrison BC, Leinwand LA. 2015. miR-30 family microRNAs regulate myogenic differentiation and provide negative feedback on the microRNA pathway. PLoS One 10. doi:10.1371/journal.pone.0118229

Guess MG, Barthel KKB, Pugach EK, Leinwand LA. 2013. Measuring microRNA reporter activity in skeletal muscle using hydrodynamic limb vein injection of plasmid DNA combined with in vivo imaging. Skelet Muscle 3:19. doi:10.1186/2044-5040-3-19

Haines CD, Harvey PA, Leinwand LA. 2012. Estrogens mediate cardiac hypertrophy in a stimulus-dependent manner. Endocrinology 153:4480–4490. doi:10.1210/en.2012-1353

Haizlip KM, Harrison BC, Leinwand LA. 2015. Sex-based differences in skeletal muscle kinetics and fiber-type composition. Physiology 30:30–39. doi:10.1152/physiol.00024.2014

Hamrick MW, Herberg S, Arounleut P, He HZ, Shiver A, Qi RQ, Zhou L, Isales CM, Mi QS. 2010. The adipokine leptin increases skeletal muscle mass and significantly alters skeletal muscle miRNA expression profile in aged mice. Biochem Biophys Res Commun 400:379–383. doi:10.1016/j.bbrc.2010.08.079

Harada K, Sugaya T, Murakami K, Yazaki Y, Komuro I. 1999. Angiotensin II type 1A receptor knockout mice display less left ventricular remodeling and improved survival after myocardial infarction. Circulation 100:2093–2099. doi:10.1161/01.CIR.100.20.2093

Harrison BC, Allen DL, Girten B, Stodieck LS, Kostenuik PJ, Bateman TA, Morony S, Lacey D, Leinwand LA. 2003. Skeletal muscle adaptations to microgravity exposure in the mouse. J Appl Physiol 95:2462–2470. doi:10.1152/japplphysiol.00603.2003

Hoyer K, Krenz M, Robbins J, Ingwall JS. 2007. Shifts in the myosin heavy chain isozymes in the mouse heart result in increased energy efficiency. J Mol Cell Cardiol 42:214–221. doi:10.1016/j.yjmcc.2006.08.116

Izumo S, Lompré AM, Matsuoka R, Koren G, Schwartz K, Nadal-Ginard B, Mahdavi V. 1987. Myosin heavy chain messenger RNA and protein isoform transitions during cardiac hypertrophy. Interaction between hemodynamic and thyroid hormone-induced signals. J Clin Invest 79:970–977. doi:10.1172/JCI112908

Janssen I, Heymsfield SB, Wang Z, Ross R. 2000. Skeletal muscle mass and distribution in 468 men and women aged 18-88 yr. J Appl Physiol 89:81–88.

Kim HK, Lee YS, Sivaprasad U, Malhotra A, Dutta A. 2006. Muscle-specific microRNA miR-206 promotes muscle differentiation. J Cell Biol 174:677–687. doi:10.1083/jcb.200603008

Kim JY, Park YK, Lee KP, Lee SM, Kang TW, Kim HJ, Dho SH, Kim SY, Kwon KS. 2014. Genome-wide profiling of the microRNA-mRNA regulatory network in skeletal muscle with aging. Aging (Albany NY) 6:524–544. doi:10.18632/aging.100677

Krenz M, Robbins J. 2004. Impact of beta-myosin heavy chain expression on cardiac function during stress. J Am Coll Cardiol 44:2390–2397. doi:10.1016/j.jacc.2004.09.044

Krenz M, Sadayappan S, Osinska HE, Henry JA, Beck S, Warshaw DM, Robbins J. 2007. Distribution and structure-function relationship of myosin heavy chain isoforms in the adult mouse heart. J Biol Chem 282:24057–24064. doi:10.1074/jbc.M704574200

Limana F, Esposito G, D’Arcangelo D, Di Carlo A, Romani S, Melillo G, Mangoni A, Bertolami C, Pompilio G, Germani A, Capogrossi MC. 2011. HMGB1 attenuates cardiac remodelling in the failing heart via enhanced cardiac regeneration and miR-206-mediated inhibition of TIMP-3. PLoS One 6:e19845. doi:10.1371/journal.pone.0019845

Liu N, Bezprozvannaya S, Shelton JM, Frisard MI, Hulver MW, McMillan RP, Wu Y, Voelker KA, Grange RW, Richardson JA, Bassel-Duby R, Olson EN. 2011. Mice lacking microRNA 133a develop dynamin 2-dependent centronuclear myopathy. J Clin Invest 121:3258–3268. doi:10.1172/JCI46267

Liu N, Williams AH, Maxeiner JM, Bezprozvannaya S, Shelton JM, Richardson JA, Bassel-Duby R, Olson EN. 2012. microRNA-206 promotes skeletal muscle regeneration and delays progression of Duchenne muscular dystrophy in mice. J Clin Invest 122:2054–2065. doi:10.1172/JCI62656

Ma G, Wang Y, Li Y, Cui L, Zhao Y, Zhao B, Li K. 2015. MiR-206, a key modulator of skeletal muscle development and disease. Int J Biol Sci 11:345–352. doi:10.7150/ijbs.10921

McCarthy JJ. 2008. MicroRNA-206: The skeletal muscle-specific myomiR. Biochim Biophys Acta - Gene Regul Mech 1779:682–691. doi:10.1016/j.bbagrm.2008.03.001

McCarthy JJ, Fox AM, Tsika GL, Gao L, Tsika RW. 1997. β-MHC transgene expression in suspended and mechanically overloaded/suspended soleus muscle of transgenic mice. Am J Physiol - Regul Integr Comp Physiol 272:R1552–61.

Mitchelson KR. 2015. Roles of the canonical myomiRs miR-1, −133 and −206 in cell development and disease. World J Biol Chem 6:162–208. doi:10.4331/wjbc.v6.i3.162

Miyata S, Minobe W, Bristow MR, Leinwand LA. 2000. Myosin Heavy Chain Isoform Expression in the Failing and Nonfailing Human Heart. Circ Res 86:386–390. doi:10.1161/01.RES.86.4.386

Nadal-Ginard B, Mahdavi V. 1989. Molecular basis of cardiac performance. Plasticity of the myocardium generated through protein isoform switches. J Clin Invest 84:1693–1700. doi:10.1172/JCI114351

Nakao K, Minobe W, Roden R, Bristow MR, Leinwand LA. 1997. Myosin heavy chain gene expression in human heart failure. J Clin Invest 100:2362–2370. doi:10.1172/JCI119776

O’Rourke JR, Georges SA, Seay HR, Tapscott SJ, McManus MT, Goldhamer DJ, Swanson MS, Harfe BD. 2007. Essential role for Dicer during skeletal muscle development. Dev Biol 311:359–368. doi:10.1016/j.ydbio.2007.08.032

Patrizio M, Musumeci M, Piccone A, Raggi C, Mattei E, Marano G. 2013. Hormonal regulation of β-myosin heavy chain expression in the mouse left ventricle. J Endocrinol 216:287–296. doi:10.1530/JOE-12-0201

Potthoff MJ, Wu H, Arnold MA, Shelton JM, Backs J, McAnally J, Richardson JA, Bassel-Duby R, Olson EN. 2007. Histone deacetylase degradation and MEF2 activation promote the formation of slow-twitch myofibers. J Clin Invest 117:2459–2467. doi:10.1172/JCI31960

Rao PK, Kumar RM, Farkhondeh M, Baskerville S, Lodish HF. 2006. Myogenic factors that regulate expression of muscle-specific microRNAs. Proc Natl Acad Sci 103:8721–8726. doi:10.1073/pnas.0602831103

Resnicow DI, Deacon JC, Warrick HM, Spudich JA, Leinwand LA. 2010. Functional diversity among a family of human skeletal muscle myosin motors. Proc Natl Acad Sci U S A 107:1053–1058. doi:10.1073/pnas.0913527107

Ross A, Leveritt M. 2001. Long-term metabolic and skeletal muscle adaptations to short-sprint training: Implications for sprint training and tapering. Sport Med 31:1063–1082. doi:10.2165/00007256-200131150-00003

Sadayappan S, Gulick J, Klevitsky R, Lorenz JN, Sargent M, Molkentin JD, Robbins J. 2009. Cardiac myosin binding protein-C phosphorylation in a β-myosin heavy chain background. Circulation 119:1253–1262. doi:10.1161/CIRCULATIONAHA.108.798983

Sakakibara I, Santolini M, Ferry A, Hakim V, Pascal M. 2014. Six Homeoproteins and a Iinc-RNA at the Fast MYH Locus Lock Fast Myofiber Terminal Phenotype. PLoS Genet 10:e1004386. doi:10.1371/journal.pgen.1004386

Shan ZX, Lin QX, Deng CY, Zhu JN, Mai LP, Liu JL, Fu YH, Liu XY, Li YX, Zhang YY, Lin SG, Yu XY. 2010. MiR-1/miR-206 regulate Hsp60 expression contributing to glucose-mediated apoptosis in cardiomyocytes. FEBS Lett 584:3592–3600. doi:10.1016/j.febslet.2010.07.027

Shan ZX, Lin QX, Fu YH, Deng CY, Zhou ZL, Zhu JN, Liu XY, Zhang YY, Li Y, Lin SG, Yu XY. 2009. Upregulated expression of miR-1/miR-206 in a rat model of myocardial infarction. Biochem Biophys Res Commun 381:597–601. doi:10.1016/j.bbrc.2009.02.097

Sweetman D, Goljanek K, Rathjen T, Oustanina S, Braun T, Dalmay T, Münsterberg A. 2008. Specific requirements of MRFs for the expression of muscle specific microRNAs, miR-1, miR-206 and miR-133. Dev Biol 321:491–499. doi:10.1016/j.ydbio.2008.06.019

Taegtmeyer H, Sen S, Vela D. 2010. Return to the fetal gene program: A suggested metabolic link to gene expression in the heart. Ann N Y Acad Sci 1188:191–198. doi:10.1111/j.1749-6632.2009.05100.x

Trexler CL, Odell AT, Jeong MY, Dowell RD, Leinwand LA. 2017. Transcriptome and Functional Profile of Cardiac Myocytes Is Influenced by Biological Sex. Circ Cardiovasc Genet 10:pii: e001770. doi:10.1161/CIRCGENETICS.117.001770

van Rooij E, Quiat D, Johnson BA, Sutherland LB, Qi X, Richardson JA, Kelm RJ, Olson EN. 2009. A family of microRNAs encoded by myosin genes governs myosin expression and muscle performance. Dev Cell 17:662–73. doi:10.1016/j.devcel.2009.10.013

Van Rooij E, Sutherland LB, Liu N, Williams AH, McAnally J, Gerard RD, Richardson JA, Olson EN. 2006. A signature pattern of stress-responsive microRNAs that can evoke cardiac hypertrophy and heart failure. Proc Natl Acad Sci U S A 103:18255–18260. doi:10.1073/pnas.0608791103

Webster C, Silberstein L, Hays AP, Blau HM. 1988. Fast muscle fibers are preferentially affected in Duchenne muscular dystrophy. Cell 52:503–513. doi:10.1016/0092-8674(88)90463-1

Westendorp B, Major JL, Nader M, Salih M, Leenen FHH, Tuana B. 2012. The E2F6 repressor activates gene expression in myocardium resulting in dilated cardiomyopathy. FASEB J 26:2569–2579. doi:10.1096/fj.11-203174 LK - http://sfx.library.uu.nl/utrecht?sid=EMBASE&issn=08926638&id=doi:10.1096%2Ffj.11-203174&atitle=The+E2F6+repressor+activates+gene+expression+in+myocardium+resulting+in+dilated+cardiomyopathy&stitle=FASEB+J.&title=FASEB+Journal&volume=26&issue=6&spage=2569&epage=2579&aulast=Westendorp&aufirst=Bart&auinit=B.&aufull=Westendorp+B.&coden=FAJOE&isbn=&pages=2569-2579&date=2012&auinit1=B&auinitm=

Williams AH, Valdez G, Moresi V, Qi X, McAnally J, Elliott JL, Bassel-Duby R, Sanes JR, Olson EN. 2009. MicroRNA-206 delays ALS progression and promotes regeneration of neuromuscular synapses in mice. Science (80-) 326:1549–1554. doi:10.1126/science.1181046

Winbanks CE, Wang B, Beyer C, Koh P, White L, Kantharidis P, Gregorevic P. 2011. TGF-β regulates miR-206 and miR-29 to control myogenic differentiation through regulation of HDAC4. J Biol Chem. doi:10.1074/jbc.M110.192625

Yang Y, Del Re DP, Nakano N, Sciarretta S, Zhai P, Park J, Sayed D, Shirakabe A, Matsushima S, Park Y, Tian B, Abdellatif M, Sadoshima J. 2015. MIR-206 Mediates YAP-Induced Cardiac Hypertrophy and Survival. Circ Res 117:891–904. doi:10.1161/CIRCRESAHA.115.306624

Yuasa K, Ando M, Nakamura A, Takeda S, Hagiwara Y, Hijikata T. 2008. MicroRNA-206 Is Highly Expressed in Newly Formed Muscle Fibers: Implications Regarding Potential for Muscle Regeneration and Maturation in Muscular Dystrophy. Cell Struct Funct. doi:10.1247/csf.08022

Yutzey KE, Konieczny SF. 1992. Different E-box regulatory sequences are functionally distinct when placed within the context of the troponin I enhancer. Nucleic Acids Res 20:5105–5113. doi:10.1093/nar/20.19.5105

Zhao Y, Samal E, Srivastava D. 2005. Serum response factor regulates a muscle-specific microRNA that targets Hand2 during cardiogenesis. Nature 436:214–220. doi:10.1038/nature03817

