## Supplementary figures and images for "miR-206 Enforces a Slow Muscle Phenotype"

### Fig 1 S1

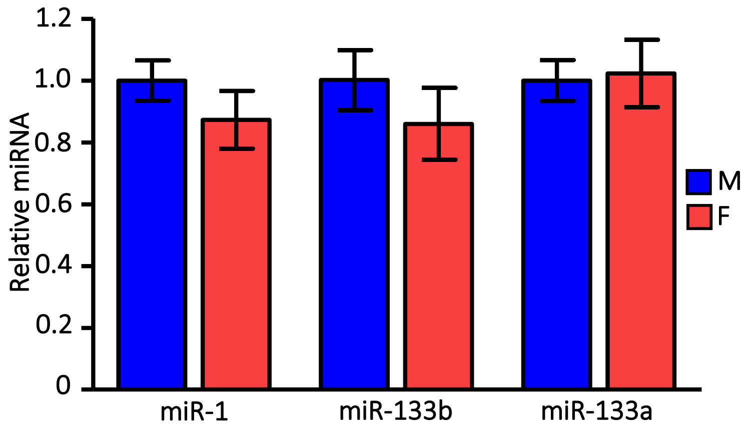

### Fig 1 S2

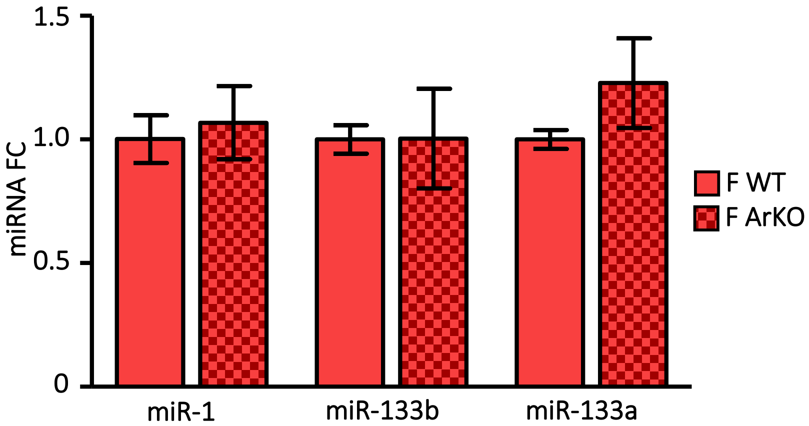

### Fig 1 S3

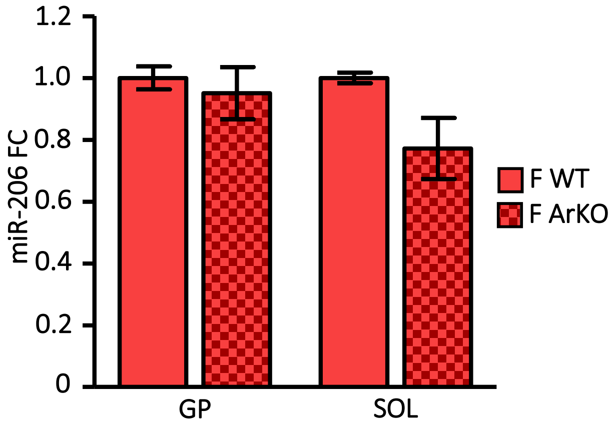

### Fig 2 S2

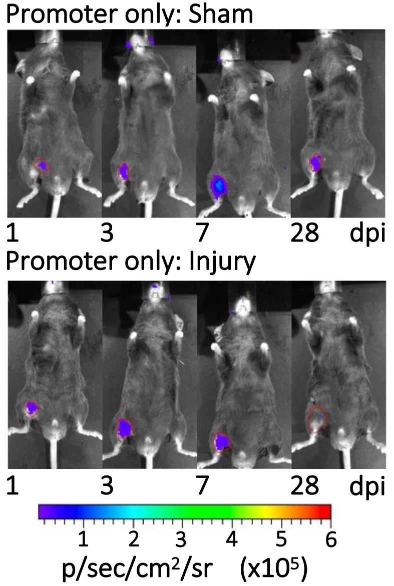

### Fig 2 S3

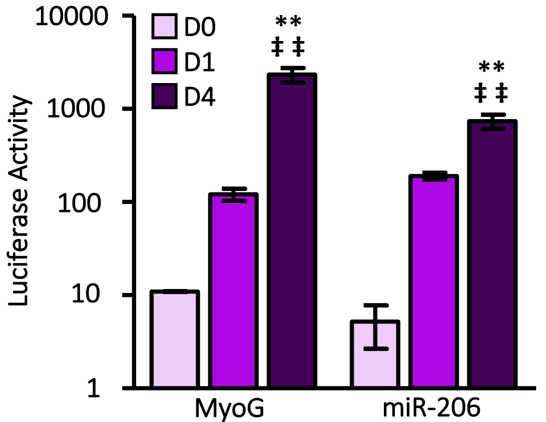

### Fig 2 S4

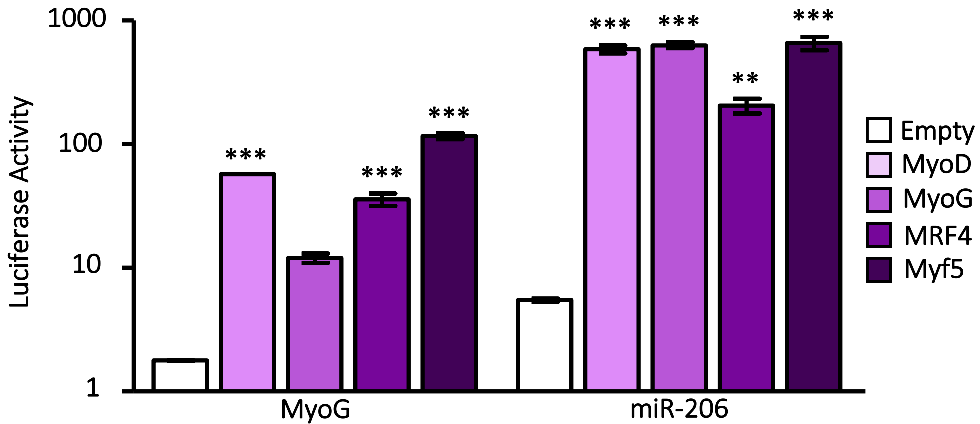

### Fig 2 S6

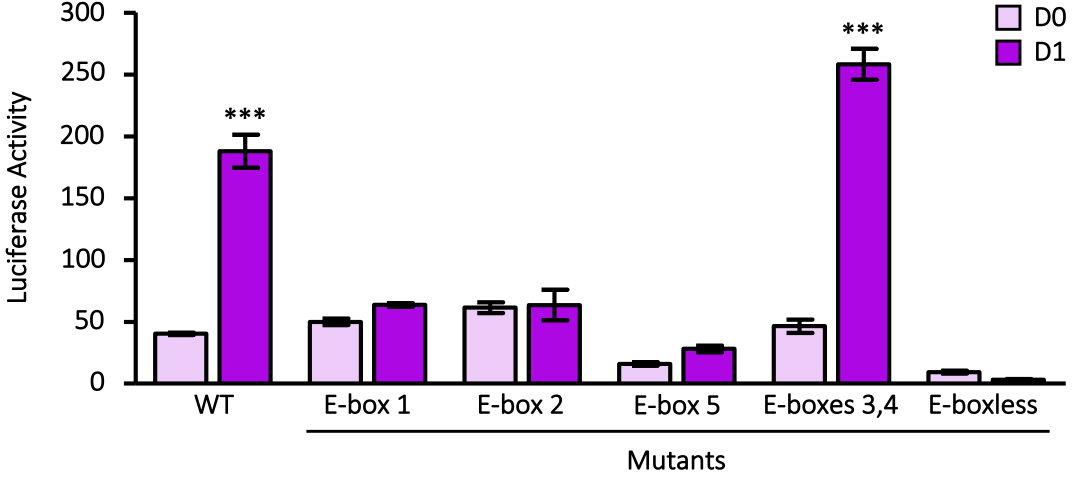

### Fig 3 S1

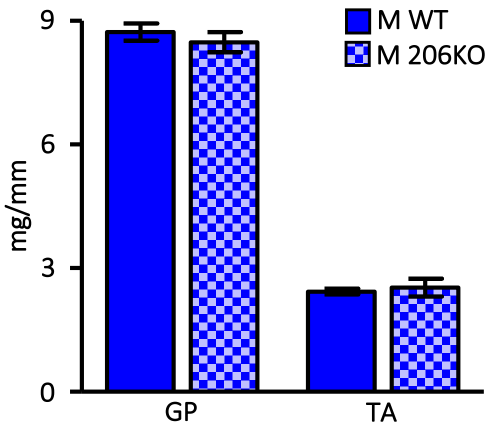

### Fig 3 S2

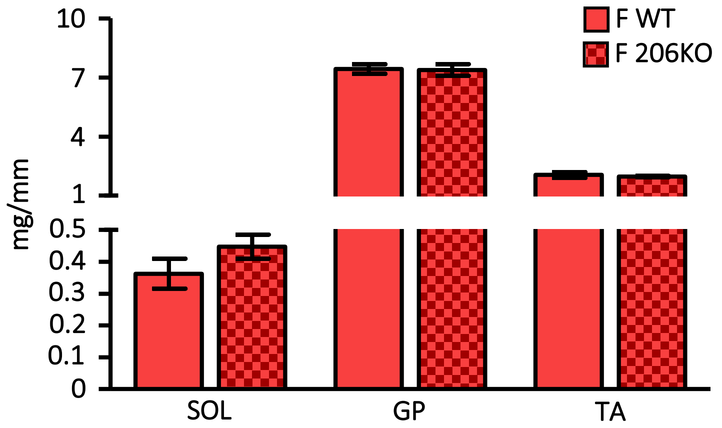

### Fig 3 S3

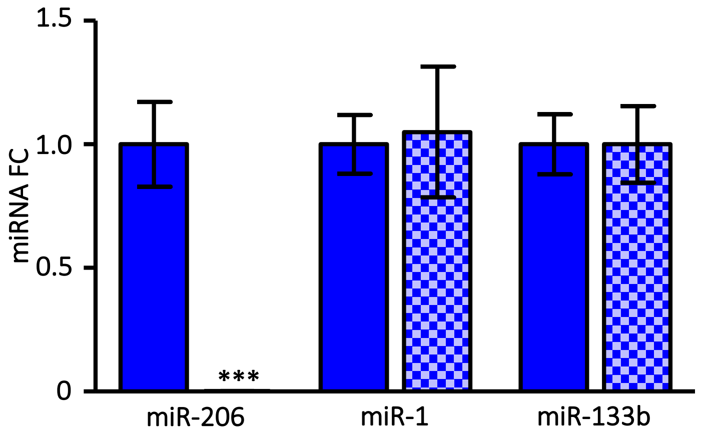

### Fig 3 S3

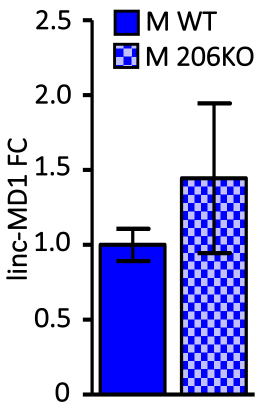

### Fig 6 S1

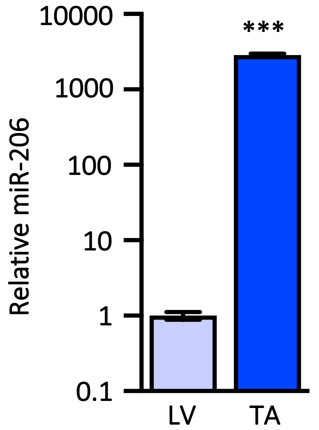

### Fig 6 S2

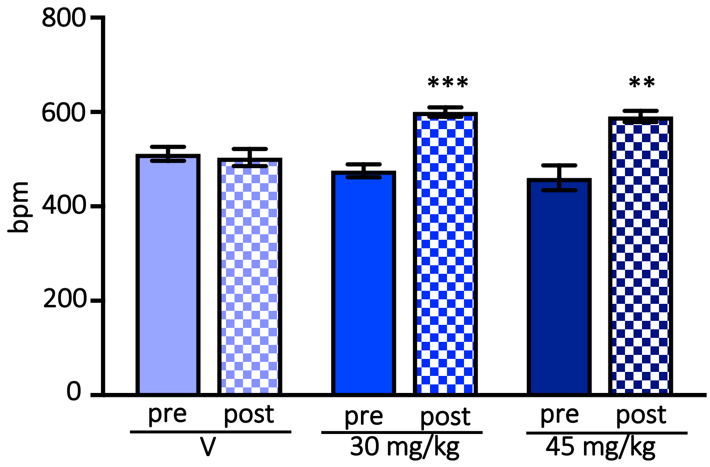

### Fig 6 S3

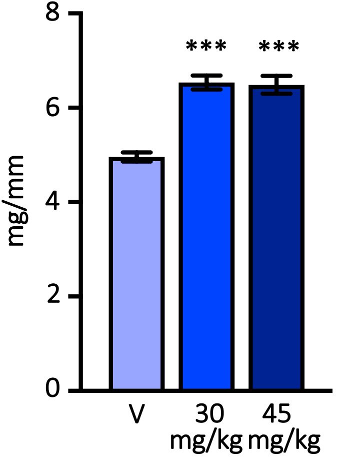

### Fig 6 S4

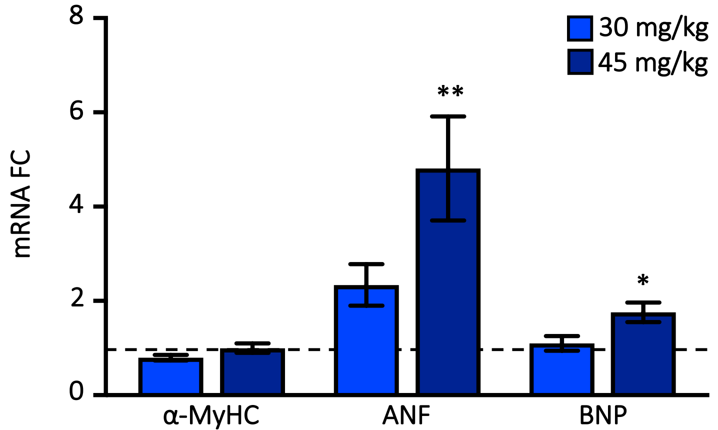

### Fig 7 S1

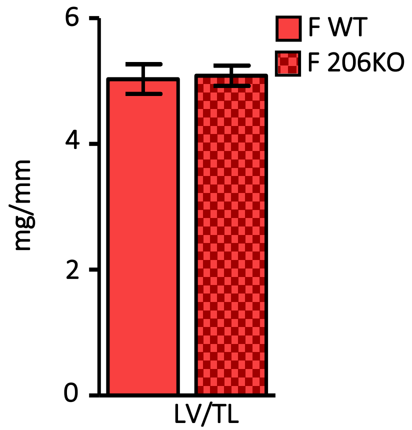

### Fig 7 S2

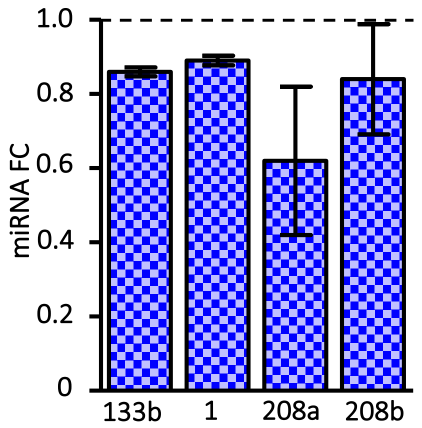

### Fig 7 S3

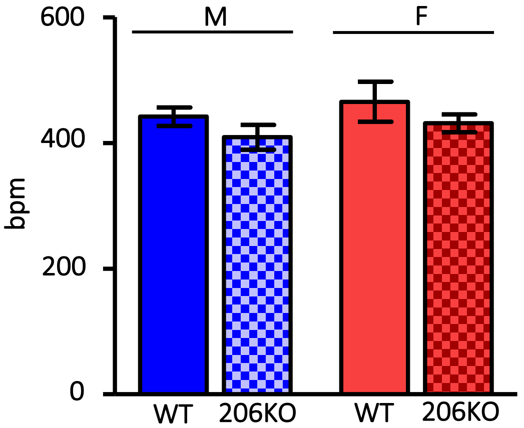

### Fig 7 S4

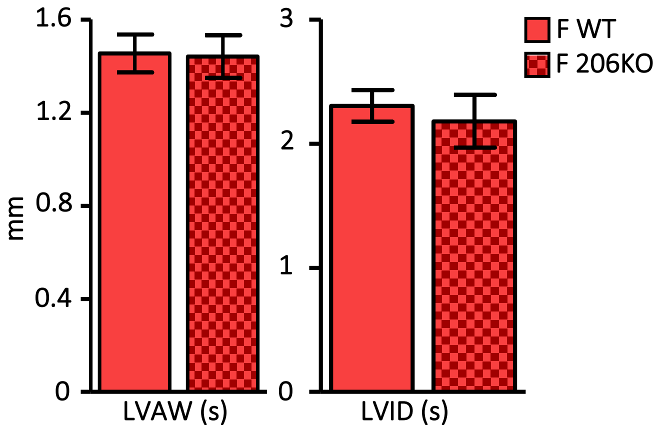

### Fig 7 S5

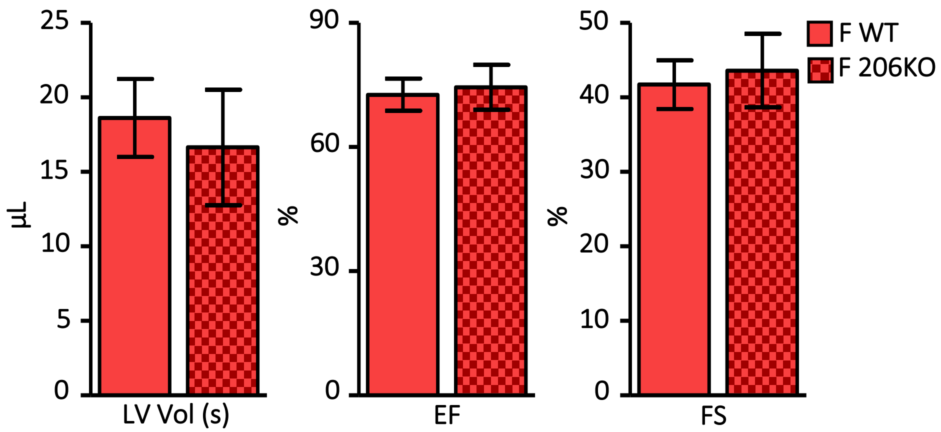

### Fig 7 S6

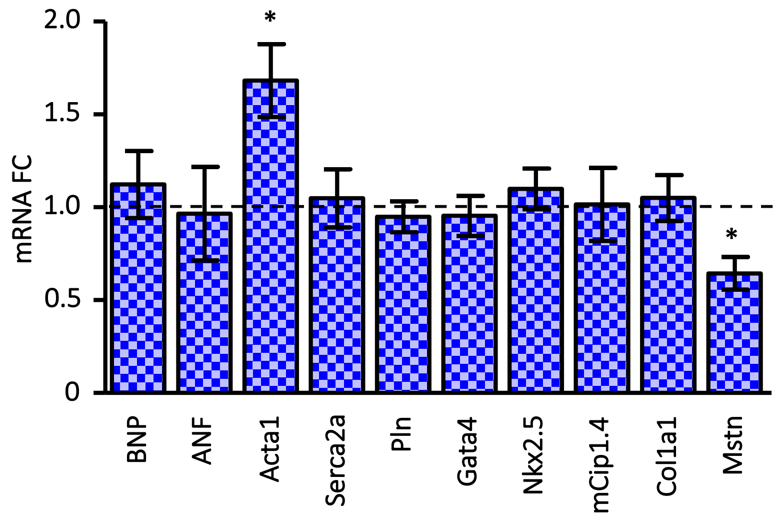

### Fig 7 S7

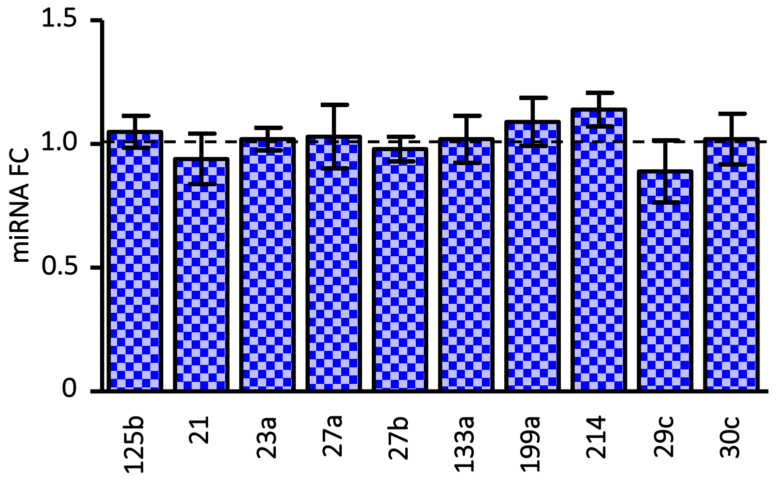
