## Supplementary material for "miR-206 Enforces a Slow Muscle Phenotype": Fig 2 S1

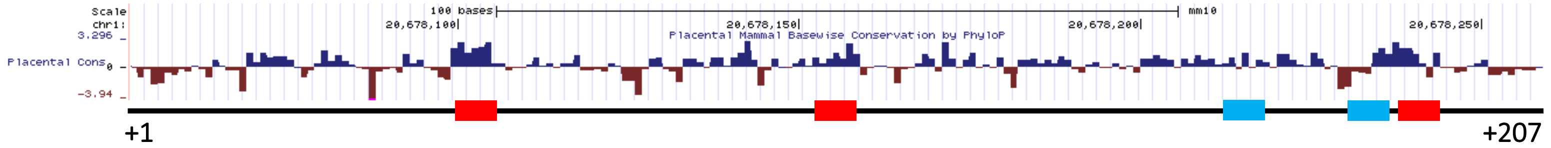


TTAATTTTCCCCATCCAGGGTGATCAGTCAGACCCTGAATGGCTCAACAGCTGCCAGTGTCCATTCATCTCCTACAGCCCAGGCCTCA
CAGATTGTGACGCAGGTGATTGGCCACGAGCAGCAGGGGCTGAACAAAAAAAGATCCCTTCCACAATGAACAATTGTGTTGGAATG
CTCATCTGGCAGCTGCAGCCTGTCAGCATATTC
