## Supplementary material for "miR-206 Enforces a Slow Muscle Phenotype": Fig 2 S5

Muscle Consensus: (C/G)N(G/A)(G/A)CA(C/G)(C/G)TG(C/T)(C/T)N(C/G)

```
E-box #   Mouse                    Score    Human                    Score
-------------------------------------------------------------------------
          *                                 *            *
  1       TCAACAGCTGCCAG  =  7.5            TCAACAGCTGCCAA  =  7.0

            *       *                                 *
  2       GACGCAGGCGATTG  =  7.0            GGGGCAGGTGATGG  =  7.5

          *     **   * *
  3       TGAACAATTGTGTT  =  5.5                  Ø

          * **  *   *
  4       TGCTCATCTGGCAG  =  5.5                  Ø

                     *                      *
  5       CTGGCAGCTGCAGC  =  7.5            TGGGCAGCTGCTGC  =  7.5
```
